# Genomic characterisation and context of the *bla*_NDM-1_ carbapenemase in *Escherichia coli* ST101

**DOI:** 10.1101/860726

**Authors:** Melinda M. Ashcroft, Brian M. Forde, Minh-Duy Phan, Kate M. Peters, Andrew Henderson, Steven J. Hancock, Leah W. Roberts, Rhys T. White, Kok-Gan Chan, Teik Min Chong, Wai-Fong Yin, David L. Paterson, Timothy R. Walsh, Mark A. Schembri, Scott A. Beatson

## Abstract

Carbapenems are last-resort antibiotics; however, the spread of plasmid-encoded carbapenemases such as the New Delhi metallo-β-lactamase 1 (NDM-1) challenges their effectiveness. The rise of NDM-1 has coincided with the emergence of extensively multidrug resistant (MDR) lineages such as *Escherichia coli* ST101. Here we present a comprehensive genomic analysis of seven *E. coli* ST101 isolates that carry the *bla*_NDM-1_ gene. We determined the complete genomes of two isolates and the draft genomes of five isolates, enabling complete resolution of the plasmid context of *bla*_NDM-1_. Comparisons with thirteen previously published ST101 genomes revealed a monophyletic lineage within the B1 phylogroup forming two clades (designated Clade 1 and Clade 2). Most Clade 1 strains are MDR, encoding resistance to at least 9 different antimicrobial classes, including extended spectrum cephalosporins. Additionally, we characterised different pathways for *bla*_NDM-1_ carriage and persistence in the ST101 lineage. For IncC plasmids, carriage was associated with recombination and local transposition events within the antibiotic resistance island. In contrast, we revealed recent transfer of a large *bla*_NDM-1_ resistance island between F-type plasmids. The complex acquisition pathways characterised here highlight the benefits of long-read Single Molecule Real Time sequencing in revealing evolutionary events that would not be apparent by short-read sequencing alone. These high-quality *E. coli* ST101 genomes will provide an important reference for further analysis of the role of mobile genetic elements in this emerging multidrug resistant lineage.

**Importance:** Carbapenem resistant *Escherichia coli* are urgent priority organisms as they are resistant to our drugs of last resort. *E. coli* ST101 have been reported as carriers of the New Delhi Metallo-beta-Lactamase 1 gene (bla_NDM-1_), conferring resistance to carbapenems, however there is limited genomic information available for this lineage. In this study we used long-read genome sequencing to characterise the complete genomes of two *E. coli* ST101 strains and determine the carriage of *bla*_NDM-1_ in a collection of *E. coli* ST101 strains. We showed that carriage of *bla*_NDM-1_ and resistance determinants to eight other antimicrobial classes was confined to a single clade. We also showed two different pathways for the carriage of *bla*_NDM-1_, which was dependent on the type of plasmid. Long-read sequencing allowed us to show the full complexities of these resistance regions and highlighted how strains from an emerging *E. coli* lineage have become resistant to nearly all available antimicrobials.

## Introduction

The successful treatment of *E. coli* infections is complicated by the rising prevalence of antibiotic resistance among clinical and community isolates (1). Carbapenems are considered a class of last-resort antibiotics, however their effectiveness has been challenged by the emergence of bacteria capable of hydrolysing carbapenems and most β-lactams. Carbapenem resistance has disseminated worldwide, as genes that encode carbapenemases are easily transferred via MGEs such as plasmids, transposons and integrons. In 2008, *bla*_NDM-1_ was first reported in a *Klebsiella pneumoniae* strain isolated from a Swedish patient who had recently travelled to India (2). Since then, the *bla*_NDM-1_ gene and its variants has been identified in Gram-negative bacteria in more than fifty countries including the UK, India, Pakistan and Bangladesh, across Europe, China, the US, Canada and Australia (3). NDM-positive clonal lineages can be found in multiple different organisms including *K. pneumoniae, Acinetobacter baumannii* and *E. coli* (4). For example, in *K. pneumoniae*, several of the first *bla*_NDM-1_ reports between 2009-2011 involved the multi-drug resistant (MDR) sequence type (ST)14 clone (5). In *A. baumannii*, international clonal lineages I (Clonal Complex (CC)109/CC1) and II (CC92/CC2), ST25 and ST85 are the four dominant NDM-positive clonal lineages (6). In *E. coli, bla*_NDM-1_ has been detected worldwide, with isolates frequently typed as ST405, ST131 and ST101 (4) and also found to carry the *bla*_CTX-M-15_ ESBL gene (7). The first characterisation of an FII plasmid carrying the *bla*_NDM-1_ gene was in the *E. coli* ST131 strain GUE (pGUE-NDM, Genbank accession: JQ364967) (8).

Other *E. coli* MDR clones such as ST131 have been well-characterised genetically (9-11), but very little is known about *E. coli* ST101 at the genomic level, despite numerous reported cases of carbapenemase-producing *E. coli* ST101 (4, 12-16). Mushtaq et al, 2011 showed that ST101 strains have acquired NDM-positive plasmids of different incompatibility groups and demonstrated diverse PFGE banding patterns for *E. coli* ST101 strains, with some evidence of clonal expansion due to clustering of pulsotypes (12). Additionally, serotyping of five ST101 isolates indicated that they were O-non-typeable:H21/42 and contained an array of virulence genes; including *fimH, pap, sfa/focDE* (fimbrial genes), *iucD, iroN* (siderophore genes), *iss, traT* (protection genes), and *tsh* (serine protease gene) (16). Other studies have reported varying levels of virulence potential in ST101 isolates; however in all cases, these strains contained numerous adhesins, autotransporters and siderophores normally associated with extra-intestinal pathogenic *E. coli* (4, 16, 17).

There are now several *E. coli* ST101 genomes and NDM-positive plasmid sequences available in public databases, but as yet there are no published analyses of any *E. coli* ST101 complete genomes. Draft assemblies provide limited information in terms of genomic context of mobile elements such as insertion sequences (IS), phages, genomic islands and plasmids. The complete assembly of these mobile elements is crucial for characterising the genomic context of resistance elements (18, 19). Here we present a comprehensive genomic analysis of *bla*_NDM-1_ carriage in *E. coli* ST101. Using a combination of Pacific Biosciences (PacBio) Single Molecule Real Time (SMRT) sequencing and Illumina sequencing, we defined the plasmid context of *bla*_NDM-1_ in seven ST101 isolates. We also report the complete genome of two representative ST101 strains: MS6192 and MS6193, including manual curation of all MGEs. Using publicly available and published ST101 draft genomes, we also examined the phylogeny of ST101 and defined two major clades (1 and 2), with the majority of Clade 1 strains encoding resistance to carbapenems and extended spectrum cephalosporins (ESCs).

## Results

### NDM-positive *E. coli* ST101 genomes

During our molecular characterisation of 16 carbapenemase-producing *E. coli* collected from India and the United Kingdom in 2008-2009 (20, 21), we found seven that were phylogroup B1, sequence type (ST)101 and NDM-positive by PCR (Supplementary Appendix and Supplementary Dataset, Table S1). To investigate the genomic context of *bla*_NDM-1_ and characterise ST101 at the genomic level we undertook whole genome sequencing (WGS). The genomes of seven NDM-positive ST101 strains were assembled from a combination of PacBio RSI or RSII long-read data and Illumina HiSeq paired-end data. In all cases we could assemble a single plasmid encoding *bla*_NDM-1_ together with several other antimicrobial resistance (AMR) genes (see below). There were sufficient long reads to enable the assembly of complete, finished quality, genome sequences for two of the strains: MS6192 and MS6193 (Supplementary Dataset, Table S2).

*E. coli* MS6192 is comprised of a single circular chromosome of 4,879,059 bp (Table 1). Four circularised contigs of 142,890 bp, 76,661 bp, 86,336 bp and 3,608 bp represent the large *bla*_NDM-1_ encoding MDR plasmid pMS6192A-NDM (4 replicons: FII and FII(pCoo) replicons and an FIA and FIB replicon, IncF replicon sequence type (IncF RST) F36/F22:A1:B20), pMS6192B (FII, F2:A-:B-), pMS6192C (Incl1, PMLST ST173) and the small cryptic plasmid pMS6192D (ColRNAI), respectively. *E. coli* MS6193 is comprised of a single circular chromosome of 4,922,872 bp. Additionally, there were three circularised contigs; 142,890 bp, 76,661 bp and 4,367 bp, which represent the large *bla*_NDM-1_ encoding MDR plasmid pMS6193A-NDM (4 replicons: FII and FII(pCoo), FIA and FIB, F36/F22:A1:B20), pMS6193B (FII, F2:A-:B-) and pMS6193C (untypeable).

**Table 1.** Summary of genomic information for *E. coli* ST101 strains MS6192 and MS6193.

|  | MS6192<br>chr<sup>1</sup> | MS6193<br>chr<sup>1</sup> | pMS6192A-NDM/<br>pMS6193A-NDM<br>(FII/FII:F1A:F1B) | pMS6192B/<br>pMS6193B<br>(FII) | pMS6192C<sup>2</sup><br>(Incl1) | pMS6193C<sup>2</sup><br>(colRNAI) | pMS6192D<br>(untypeable) |
| --- | --- | --- | --- | --- | --- | --- | --- |
| Size<br>(bp) | 4,879,059 | 4,922,872 | 142,890/<br>142,890 | 76,661/<br>76,661 | 86,336 | 4,367 | 3,608 |
| No. of<br>genes | 4,834 | 4,876 | 179/173 | 91/84 | 95 | 6 | 2 |
| No. of<br>CDS | 4,570 | 4,612 | 169/167 | 82/82 | 91 | 5 | 1 |
| No. of<br>tRNAs | 94 | 94 | - | - | - | - | - |
| No. of<br>rRNAs | 22 (7:16S,<br>7:23S,<br>8:5S) | 22 (7:16S,<br>7:23S,<br>8:5S) | - | - | - | - | - |
| No. of<br>IS | 46 | 44 | 21/21 | 5/5 | 0 | 0 | 0 |
| G+C | 50.69 | 50.69 | 51.5/51.50 | 51.91/51.91 | 50.14 | 54.39 | 43.15 |
| content (%) |  |  |  |  |  |  |  |

<sup>1</sup>chr = chromosome. <sup>2</sup>MS6192 has 4 plasmids (pMS6192A, pMS6192B, pMS6192C and pMS6192D); MS6193 has 3 plasmids (pMS6193A, pMS6193B and pMS6193C); pMS6192A-NDM and pMS6192B are near identical to pMS6193A-NDM and pMS6193B, respectively; there is no genetic relationship between pMS6192C, pMS6193C or pMS6193D.

The chromosomes of MS6192 and MS6193 were almost identical, differing by 12 single nucleotide polymorphisms (SNPs). The sequence of the F-type (bla_NDM-1_) and FII plasmids were also near-identical between strains. Further investigation of the automated assembly was required to unambiguously determine the sequences of pMS6192A-NDM and pMS6192C, primarily due to shared regions of high nucleotide identity (Supplementary Appendix). The genomes could also be distinguished by the presence of an additional prophage (Phi8) and Transposon 7 (Tn7)-like Transposon in *E. coli* MS6193 (Supplementary Appendix, Fig S1), an additional large plasmid in MS6192 (pMS6192C) and two different cryptic plasmids (pMS6192D and pMS6193C).

MS6194, MS6201, MS6203, MS6204 and MS6207 strains were assembled as draft genomes with gapped chromosomes and circularised plasmid sequences. The five draft *E. coli* ST101 genomes range in size from 5,150,883 to 5,434,929 bp with the number of contigs ranging from 16 to 55 (Supplementary Dataset, Table S2). Each draft genome contained a single *bla*_NDM-1_ encoding MDR plasmid and several plasmid replicons ranging from one (MS6203) to seven (MS6204).

### Phylogenomic analysis of *E. coli* ST101 reveals a monophyletic lineage within the B1 phylogroup

To explore the context of ST101 within the *E. coli* phylogeny we carried out a phylogenetic reconstruction of 65 representative, publicly available, complete *E. coli* genomes together with 20 ST101 genomes. These included the seven ST101 genomes sequenced in the present study and 13 *E. coli* ST101 draft genomes that were previously published and available in GenBank or the Sequence Read Archive (SRA) on October 1^st^, 2016 (Supplementary Dataset, Table S3). The additional ST101 genomes were obtained from several clinical sources and included isolates from urine (n = 5), blood (n = 4), wound (n=1) and intestinal microbiota (n = 2). A single environmental isolate from untreated wastewater was also included. All 20 *E. coli* ST101 strains cluster together in a single lineage within the B1 phylogroup (Figure 1A). Based on the representative complete genomes included in this phylogeny, the *E. coli* ST101 lineage is most closely related to *E. coli* SE11, a fecal strain of serotype O152:H28, isolated from a healthy adult (22).

**Figure 1.**
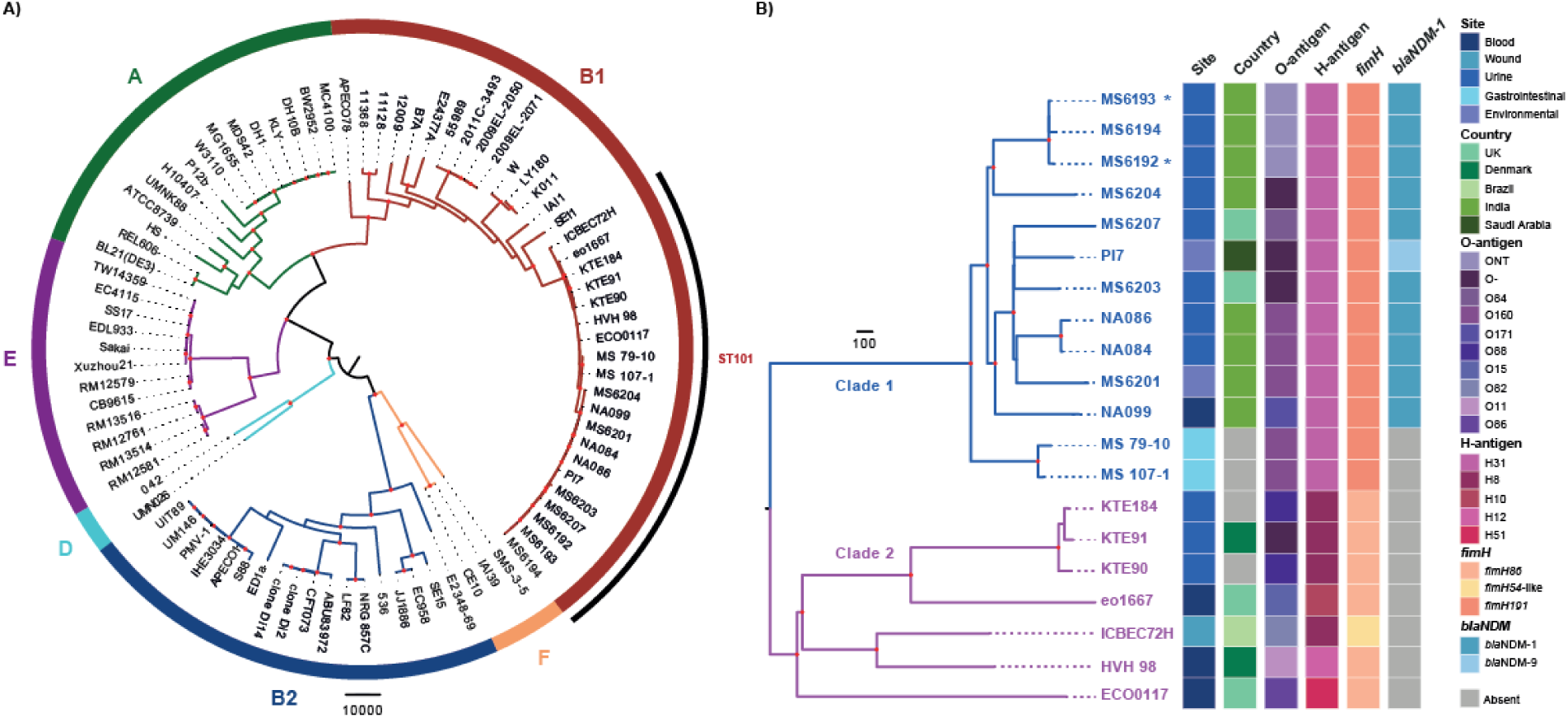
Phylogenetic relationship of *E. coli* ST101. A) The phylogram was built from 182,264 core-genome SNPs using Maximum Likelihood. The tree was rooted using the out-group species *E. fergusonii* ATCC35469 (not shown). The taxa labels for complete *E. coli* genomes are coloured as follows; phylogroup F: orange, phylogroup B2: dark blue, phylogroup D: aqua, phylogroup E: purple, phylogroup A: green, phylogroup B1: red (ST101 labelled). B) The mid-point rooted, recombination-filtered phylogram was built from 2,106 core-genome SNPs. Two clades can be defined. Clade 1 (blue): MS6193*, MS6194, MS6192*, MS6204, MS6207, PI7, MS6203, NA086, NA084, MS6201, NA099, MS 79-10 and MS 107-1; Clade 2 (pink): KTE184, KTE91, KTE90, eo1667, ICBEC72H, HVH 98 and ECO0117. NT: non-typeable, - indicates absence of locus. Bootstrapping support indicated by red nodes: 95-100% bootstrap support from 1000 replicates. *Denotes complete genomes.

The ST101 phylogeny formed two distinct clades, with Clade 1 showing more recent clonal expansion compared with the deeper branching Clade 2 (Fig 1B). Clade 1 is mostly comprised of urine isolates and includes all NDM-positive ST101, including the seven ST101 isolates sequenced in this study. In fact, only the two gut-derived isolates (MS 107-1 and MS 79-10) lack the *bla*_NDM-1_ gene (or its single amino acid variant *bla*_NDM-9_ (E152K), present in PI7). The phylogenetic position of MS 107-1 and MS 79-10, basal to the remaining NDM-positive ST101 strains, suggests that *bla*_NDM-1_ was acquired once in Clade 1 and has been inherited vertically.

To further characterise the ST101 lineage we examined key virulence and serotyping gene loci. Clade 1 isolates have the *fimH*191 type 1 fimbrial adhesion allele, whereas Clade 2 isolates possess the *fimH*86 allele except for ICBEC72H, which encodes a novel *fimH*54-like variant. Several serogroups were predicted across both clades: O160 (MS107-1, MS79-10, MS6201, NA084 and NA086), O171 (NA099), O84 (MS6207) and a novel, non-typeable O antigen region (MS6192, MS6193 and MS6194), but Clade 1 is distinguished by a *fliC* gene encoding the H31 antigen. Additionally, three Clade 1 strains lack part of the O antigen gene locus: MS6203, MS6204 and PI7 (*wzx, wzy, wzm* and *wzt*).

### Mobile genetic elements are variably conserved in *E. coli* ST101

The major differences between these ST101 genomes were MGEs, such as those defined in *E. coli* MS6192 and MS6193 (Figure 2 and Supplementary dataset, Table S4). MGEs in MS6192 and MS6193 include genomic islands (GI) within known integration tRNA hotspots (GI-*leuX*, GI-*pheU*, GI-*pheV*, GI-*glyU*), prophage elements and a Transposon-7 (Tn7)-like transposon (Supplementary Dataset, Table S5). While no MGE that was defined in MS6192 or MS6193 is completely conserved across the ST101 lineage, GI-*leuX* is present in most strains from Clades 1 and 2, and Prophage (Phi) 3 is conserved in all Clade 1 strains. MS6192, MS6193 (and MS6194) differ in only two chromosomal MGEs (Phi8 and Tn7-like Tn) consistent with their near identical core chromosomes. In contrast to the GI-*pheV* islands of phylogroup B2 ExPEC/UPEC strains, such as the 75,054 bp locus in the *E. coli* ST131 strain EC958 (23, 24), the GI within the tRNA-*pheV* locus in MS6192 and MS6193 is relatively small (9,191 bp) and does not contain known virulence modules. Other genomic regions of difference (RD) that are known to be variable across *E. coli* are present in MS6192 and MS6193 (Supplementary Dataset, Table S6) and are also conserved across the ST101 lineage. These include a remnant of the Type VI secretion system (T6SS) (25), the Flag-2 lateral flagellar locus (26) and a degenerate Type III secretion locus 2 (ETT2), most commonly found in intestinal pathogenic *E. coli* from the B1 phylogroup and rarely found in ExPEC (27). *E. coli* ST101 strains MS6192 and MS6193 also share 44 chromosomal insertion sequences (IS) (Supplementary Dataset, Table S7).

**Figure 2.**
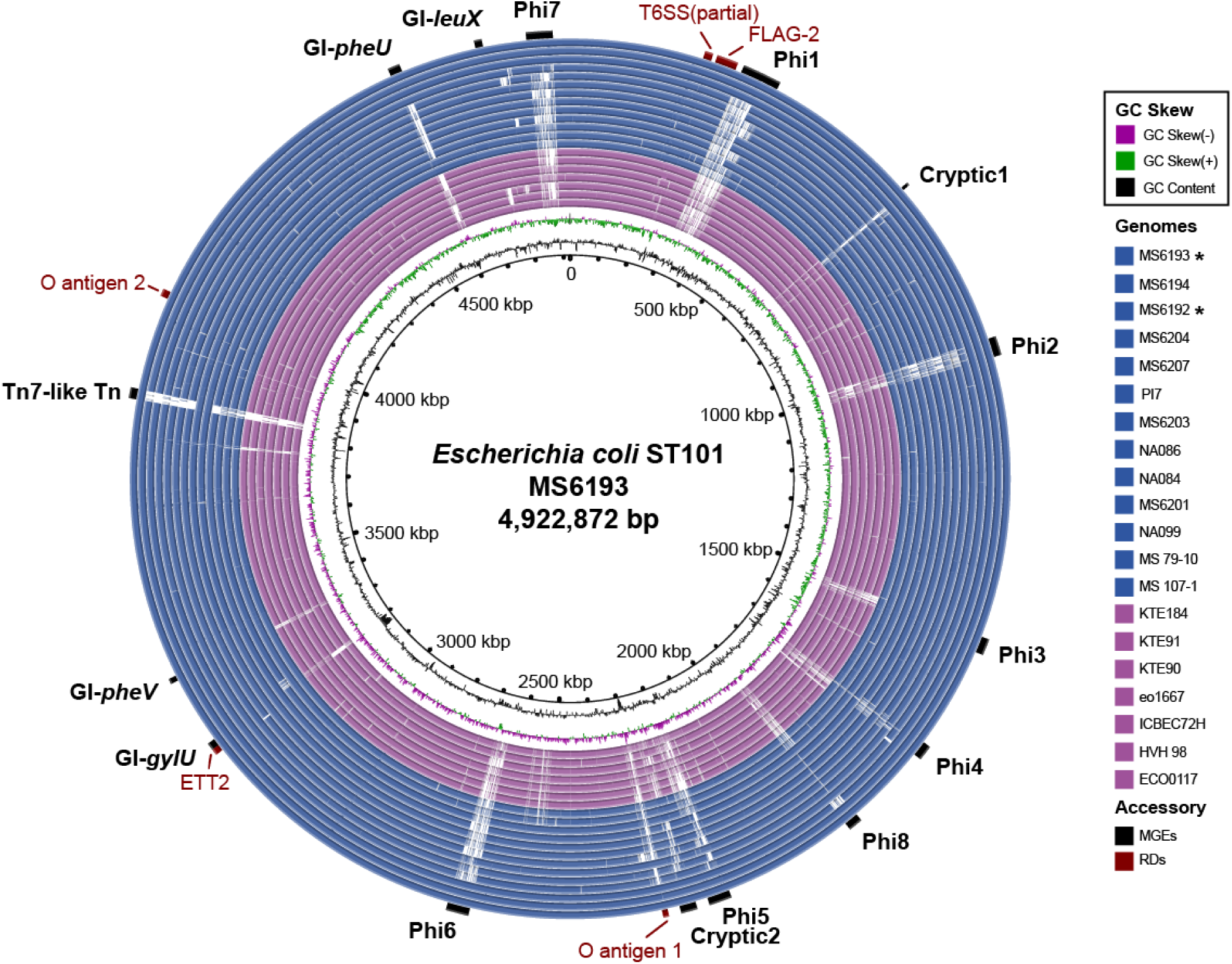
Genomic map of ST101 *E. coli* showing mobile genetic elements and other notable genomic regions. Visualisation of the *E. coli* MS6193 genome compared to 19 *E. coli* ST101 complete and draft genomes. The innermost circles represent GC skew (green/purple) and GC content (black) of *E. coli* MS6193. The degree of coloured shading indicates the nucleotide identity between MS6193 and each *E. coli* ST101 genome. Nucleotide comparisons are coloured based on identity of between 70% and 100% (dark shading = high, light shading = low). *E. coli* genomes are arranged according to their previously defined phylogenetic relationship as follows from the innermost ring: Clade 2 (pink): ECO0117, HVH 98, ICBEC72H, eo1667, KTE90, KTE91, KTE184; Clade 1 (blue): MS 107-1, MS 79-10, NA099, MS6201, NA084, NA086, MS6203, PI7, MS6207, MS6204, MS6192*, MS6194, MS6193*. *Denotes complete genomes. Outer ring indicates mobile genetic elements (MGEs) and select ST101 regions of difference (RDs) within the *E. coli* MS6193 genome. Image prepared using BRIG.

### Most *E. coli* ST101 Clade 1 strains contain multiple antibiotic resistance genes

ST101 strains from Clade 1 possessed a high number of antibiotic resistance genes, including genes that encode resistance to beta-lactams (including carbapenems), aminoglycosides, streptomycins, phenicols, sulphonamides, macrolides, trimethoprims and tetracyclines (Figure 3). All NDM-positive ST101 Clade 1 isolates also contain at least one copy of the *bla*_CTX-M-15_ gene that confers resistance to cephalosporins. In other *E. coli, bla*_CTX-M-15_ is often found on conjugative F-type plasmids as part of a Tn3-like IS*Ecp1-bla*_CTX-M-15_-orf477 mobile element (11). We could determine the genomic location of *bla*_CTX-M-15_ in all seven of the PacBio sequenced ST101 strains, showing that the *bla*_CTX-M-15_ insertion site was chromosomally located in six of the seven strains (Supplementary Dataset, Table S8). It appears that *bla*_CTX-M-15_ has been acquired once and mobilised to several different genomic locations by an IS*Ecp1* in Clade 1 and inherited vertically in some cases (Supplementary Appendix).

**Figure 3.**
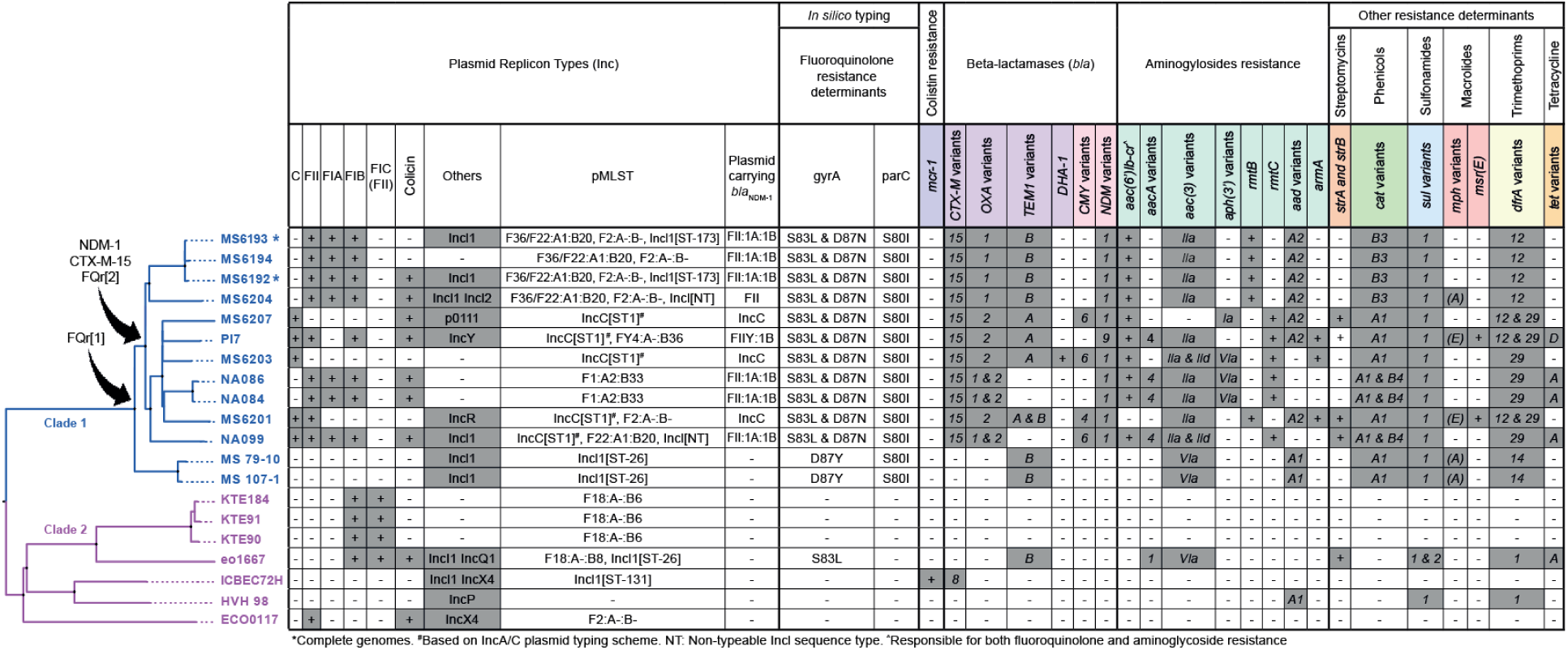
Plasmid and antimicrobial resistance typing in *E. coli* ST101. Plasmid replicon types, in silico typing and antimicrobial resistance determinants indicate similar patterns to the core-genome phylogeny (left) of the 20 *E. coli* ST101 strains. *Bla*ck arrows indicate acquisition of fluoroquinolone resistance ([1] indicates single mutation in *gyrA*, [2] indicates double mutation), *bla*NDM-1 and *bla*CTX-M-15 genes. Types of resistance genes are colour coded: colistin resistance (blue-purple) beta-lactamases (pink-purple), carbapenemases (pink), aminoglycosides (aqua), streptomycins (orange), phenicols (green), sulphonamides (blue), macrolides (red), trimethoprims (yellow) and tetracylins (gold). Grey shading indicates gene presence, white shading indicates absence, allele variants are indicated in each cell.

Fluoroquinolone resistance in *E. coli* is frequently associated with chromosomal point mutations in the quinolone resistance-determining region (QRDR) of *gyrA* (codons 67-106) and *parC* (codons 56-108) (28). All Clade 1 strains had a non-synonymous mutation in the QRDR of *parC* (S80I) (Figure 3). Additionally, two non-synonymous mutations in the QRDR of *gyrA* were detected in all NDM-positive Clade 1 isolates, characterised by the amino acid substitutions S83L and D87N. The two Clade 1 intestinal isolates MS107-1 and MS79-10 have a different single amino acid substitution in *gyrA* (D87Y) but lack both *bla*_NDM-1_ and *bla*_CTX-M-15_. These results suggest recent development of extensive MDR in a sub-lineage of Clade 1 (hereafter referred to as Clade 1A), similar to the emergence of other fluoroquinolone MDR resistant *E. coli* clonal lineages such as ST131 and ST1193 (9, 29). Phenotypic antimicrobial resistance testing of the seven PacBio sequenced ST101 strains in this study confirmed that phenotype could be predicted from genotype in most cases (Supplementary Appendix and Supplementary Dataset, Table S9). Notably Clade 2 strains did not encode QRDR mutations, with only HVH 98, ICBEC72H and eo1667 possessing any acquired resistance genes.

### The *bla*_NDM-1_ genetic environment is highly conserved in *E. coli* ST101

All seven PacBio sequenced strains contained a plasmid-encoded *bla*_NDM-1_ gene (Figure 4). In almost all cases the genetic structure of the *bla*_NDM-1_ module was identical and matched other previously described *bla*_NDM-1_ modules, consisting of a 258-287 bp fragment of ISAba125 containing the -35 promoter region, the *bla*_NDM-1_ gene, the bleomycin resistance gene (*ble*_MBL_) and a truncated phosphoribosylanthrantile isomerase (*iso/trpC*) (30). The surrounding plasmid context differed between the seven strains, but like *bla*_CTX-M-15_, the structural variations (inversions, transpositions, and insertions/deletions) are congruent with a single *bla*_NDM-1_ acquisition prior to divergence of Clade 1A. For example, only pMS6203A-NDM has retained a complete IS*Aba125* element upstream of *bla*_NDM-1_ suggestive of horizontal transfer. Additionally, downstream of the *bla*_NDM-1_ module, *iso/tat* has been truncated by the insertion of the *bla*_DHA-1_ gene. In pMS6201A-NDM, truncation of IS*Aba125* was mediated by the insertion of an IS*3000*. The two IS*3000* flanking the *bla*_NDM-1_/*ble*_MBL_ region form the composite transposon Tn*3000* (31). Truncation of ISAba125 in pMS6192A-NDM, pMS6193A-NDM, pMS6194A-NDM, pMS6204A-NDM and pMS6207A-NDM appears to have resulted from the insertion of an IS*Ecp1* module, which was subsequently deleted by an IS*26* composite transposon leaving only a 103 bp fragment of IS*Ecp1*. However, pMS6207A differs from the other four strains with carriage of *aphA6* (amikacin resistance) instead of *aac(3’)-lla* and *tmrB* (aminoglycoside and tunicamycin resistance) on the IS*26* composite transposon.

**Figure 4.**
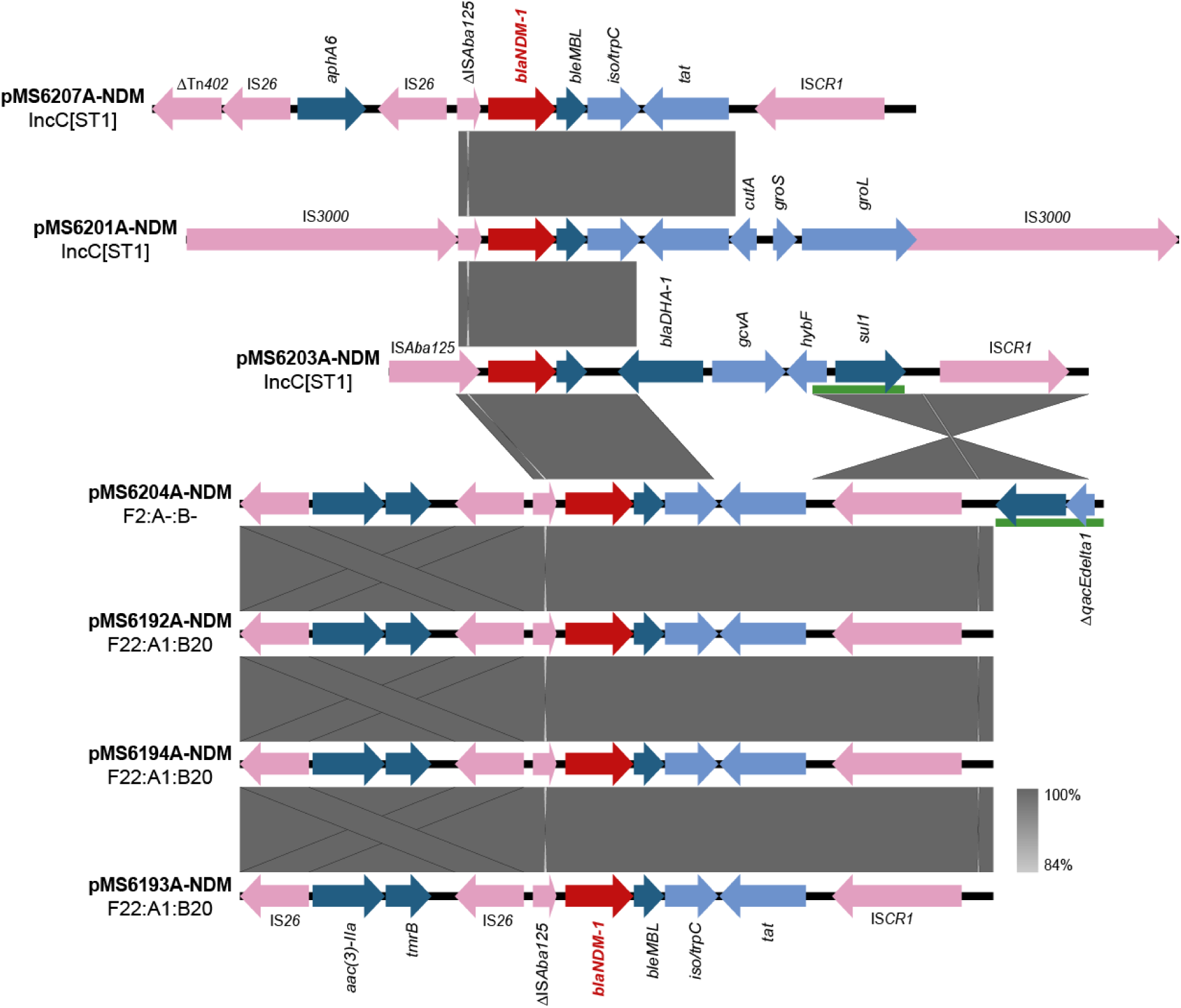
Comparison of the *bla*_NDM-1_ genetic environment of PacBio sequenced Clade 1 ST101 isolates. Grey shading indicates nucleotide identity between sequences according to *BLA*STn (84-100%). pMS6204A has been reverse complemented for easier visualisation. Δ, truncated gene. IS/Tns: light pink, AMR genes: teal, CDSs: light blue. Green rectangles represent 3’ conserved sequences of class 1 integrons. The *bla*_NDM-1_ gene is labelled in red. Image created using EasyFig.

### IncC plasmids are a vehicle for *bla*_NDM-1_ carriage in *E. coli* ST101

IncC plasmids are commonly reported carriers of the *bla*_NDM-1_ gene in *Enterobacteriaceae* (32-34). Three PacBio sequenced ST101 strains from this study contained IncC replicons (pMS6201A-NDM, pMS6203A-NDM and pMS6207A-NDM) and possess a similar backbone compared to other IncC plasmids such as pNDM-US-2 (Supplementary Appendix, Fig S2). However, deletions in the IncC plasmid backbone alter the conjugation efficiency of these plasmids such that only pMS6203A is conjugative (Supplementary Appendix). *In silico* plasmid multi-locus sequence typing (pMLST) determined that the IncC plasmids reported here belong to the NDM-associated ST1 group (34), with pMS6203A and pMS6207A belonging to the cgST 1.5. Plasmid pMS6201A is missing several genes, however, is most like cgST 1.1 (Supplementary Appendix). These IncC plasmids differ greatly in their structure of the antibiotic resistance island (ARI-A) (Supplementary Appendix and Figure 5). Therefore, despite high conservation in the immediate genetic context of *bla*NDM-1 and the IncC plasmid backbone in these ST101 strains, the complete plasmid sequences reveal substantial diversity within the ARI-A, consistent with mobilisation of AMR genes, including *bla*_NDM-1_.

**Figure 5.**
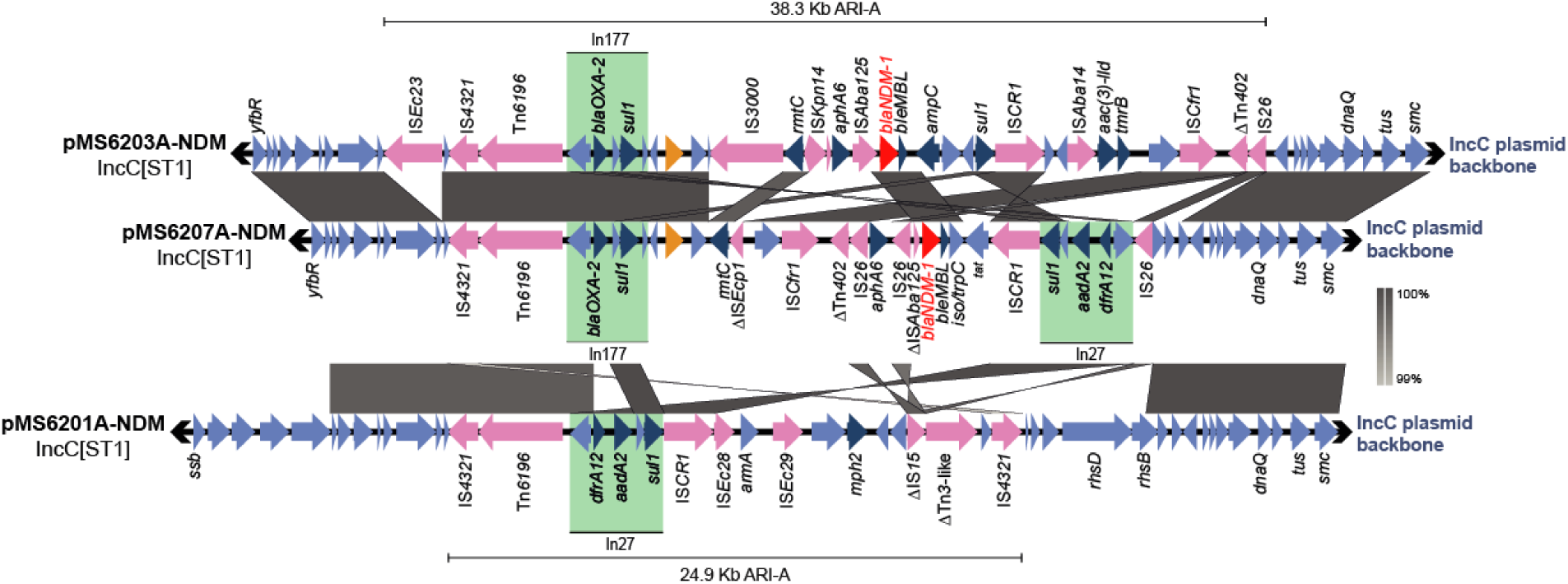
Major structural features of the Antibiotic Resistance Island on IncC plasmids in the PacBio sequenced Clade 1 ST101 strains. Nucleotide comparisons between the variable resistance regions of the IncC plasmid pMS6203A-NDM, pMS6207A-NDM and pMS6201A-NDM. Grey shading indicates nucleotide identity between sequences according to *BLA*STn (98-100%). Key genomic regions are indicated: IS elements/transposons: light pink, replication genes: dark pink, maintenance genes: purple, *pemIK* operon: aqua, MTases: orange, AMR genes: teal, other CDSs: blue, Integrons: green rectangles. The *bla*NDM-1 gene is labelled in red. Image created using EasyFig.

### *E. coli* ST101 strains possess distinct types of *bla*_NDM-1_ containing F-type plasmids

F-type plasmids have previously been associated with the carriage of *bla*_NDM-1_ in *E. coli* (8, 35-37). The four *bla*_NDM-1_ encoding F-type plasmids sequenced in this study share a near identical 21.6 kb MDR region that includes the *bla*_NDM-1_ gene. However, two different plasmid backbones were identified; plasmids pMS6192A-NDM, pMS6193A-NDM, pMS6194A-NDM were almost identical, whereas the backbone of pMS6204A-NDM differed substantially (Supplementary Appendix, Fig S3). *BLA*ST comparisons of the *bla*_NDM-1_ containing plasmids pMS6192A-NDM, pMS6193A-NDM and pMS6194A-NDM (all F36/F22:A1:B20), indicate that they are most similar to the F-type plasmid pIP1206 (>99% nucleotide identity, 83% query coverage). Major differences include deletions in the plasmid backbone, where pMS6192A-NDM, pMS6193A-NDM and pMS6194A-NDM contain an incomplete conjugation region, missing *traK* and *traB*, as well as an 18,516 bp section between *traU* and *traI* present in pIP1206. Resistance genes in pMS6192A-NDM, pMS6193A-NDM and pMS6194A-NDM are clustered in a 21.6 Kb resistance-island, which is different to the resistance island of pIP1206 and is highly similar (99.99% nucleotide identity, 91% query coverage) to the resistance island in the FII plasmid pGUE-NDM (Supplementary Appendix, Fig S4).

The *bla*_NDM-1_ resistance island has inserted between the *pemIK* toxin/antitoxin operon and the scsCD operon (copper-sensitivity suppressor). It is composed of a Tn*5403* transposon (containing an intact transposase and the inverted left repeat (ILR) (38)) truncated by an IS*26* (Figure 6). This is followed by 3 resistance genes; *aac(6’)Ib-cr* (fluoroquinolone and aminoglycoside resistance), *bla*_OXA-1_ (beta-lactam resistance) and a truncated *catB3* (chloramphenicol resistance). Two additional resistance genes *aac(3)-IIa* (aminoglycoside resistance) and *tmrB* (tunicamycin resistance) are downstream and flanked on either side by IS*26* elements. The *bla*_NDM-1_ module consists of an IS*Aba125* fragment containing the *bla*_NDM-1_ -35 promoter region, *bla*_NDM-1_ (carbapenem resistance), ble_MBL_ (bleomycin resistance), *iso/trpC* (phosphoribosylanthranilate isomerase), *tat* (twin-arginine translocation pathway signal protein) and an IS*CR1* element. Downstream of the *bla*_NDM-1_ module is the Class I integron In27. This contains *sul1* (sulphonamide resistance) and a truncated *qacEdelta1*, followed by *aadA2* (aminoglycoside resistance) and *dfrA12* (trimethoprim resistance). This MDR region ends with a truncated Tn*As3* transposon (containing intact transposase and recombinase genes and the inverted right repeat (IRR) (38, 39)).

**Figure 6.**
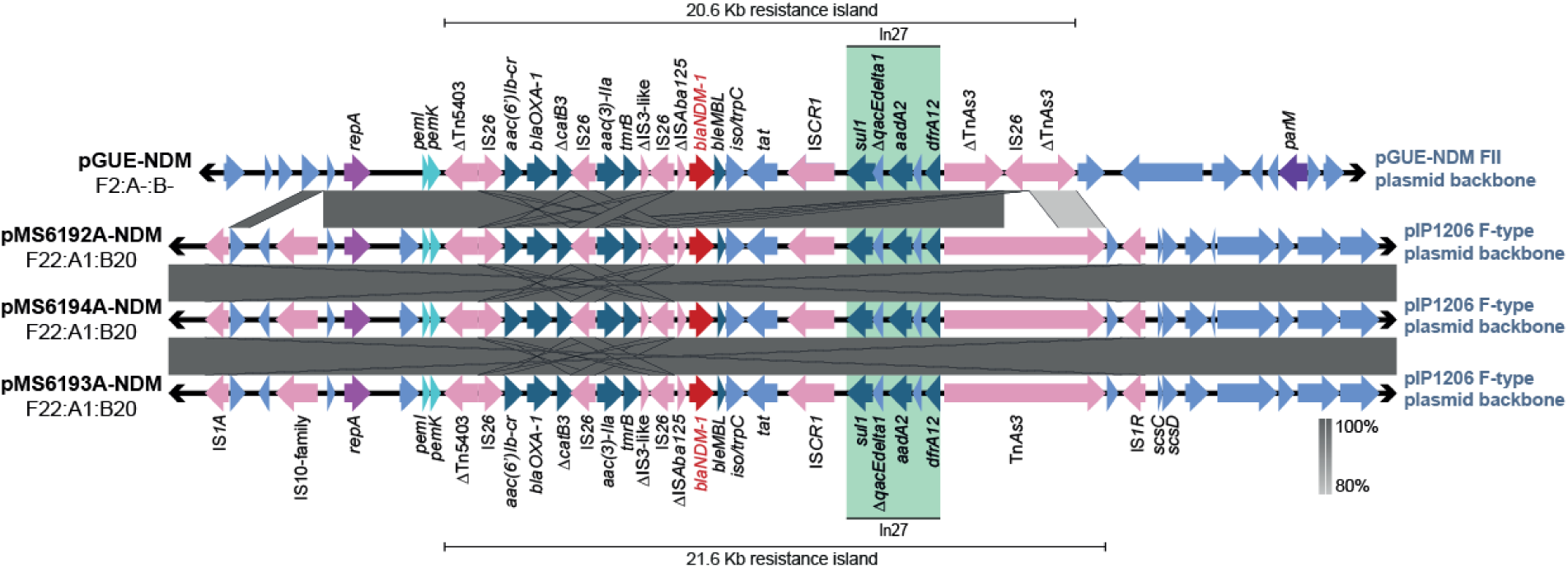
Major structural features of the conserved MDR island on F-type plasmids in the PacBio sequenced Clade 1 ST101 strains. Nucleotide comparisons between the reference plasmid pGUE-NDM (Genbank accession: JQ364967), pMS6192A-NDM, pMS6194A-NDM and pMS6193A-NDM highlighting the shared resistance island and differing plasmid backbones. pGUE-NDM has been reverse complemented for easier visualisation. Grey shading indicates nucleotide identity between sequences according to *BLA*STn (80-100%). Key genomic regions are indicated: IS elements/transposons: light pink, replication genes: dark pink, maintenance genes: purple, *pemIK* operon: aqua, MTases: orange, AMR genes: teal, other CDSs: blue, Integrons: green rectangles. The *bla*NDM-1 gene is labelled in red. Image created using EasyFig.

### bla_NDM-1_ is encoded on an FII, pGUE-NDM-like plasmid in one ST101 strain

MS6204 also contains an F-type pIP1206-like plasmid (pMS6204B; F36/F22:A1:B20). However, the resistance island in pMS6204B is different to that of pIP1206 and to that of the other *bla*_NDM-1_ containing plasmids described above. This resistance island does not carry the *bla*_NDM-1_ gene, however does encode *bla*_CTX-M-15_ and additional copies of the *aac(6’)-Ib-cr, bla*_OXA-1_ and tmrB resistance genes. In fact, for MS6204, the MDR region encoding *bla*_NDM-1_ is harboured by pMS6204A-NDM (F2:A-:B-), an FII plasmid that is most similar to *E. coli* FII plasmid pGUE-NDM (>99% nucleotide identity, 95% query coverage). Additionally, present within the MDR region of the pMS6204A-NDM plasmid is a 5,095 bp resistance region, which is not present in pGUE-NDM. This region is flanked on either side by IS26 and encompasses a Tn3 transposon encoding *bla*_TEM-1B_ (beta-lactam resistance) followed by *rmtB* (rifampicin resistance). This association between Tn*3, bla*_TEM-1_ and *rmtB* flanked by IS*26* has been observed in several other *E. coli* and *K. pneumoniae* plasmids (Supplementary Dataset, Table S10), suggesting widespread distribution of this resistance module.

### Complete plasmid sequences from closely related ST101 strains show carriage of *bla*_NDM-1_ on different F-type plasmids

Intriguingly MS6192, MS6193 and MS6194 also contain a pGUE-NDM-like plasmid, with almost identical backbones to pMS6204A-NDM (F2:A-:B-; named pMS6192B and pMS6193B in our complete genomes), however, these plasmids lack the *bla*_NDM-1_ encoding MDR island (Supplementary Appendix, Fig S4). We therefore hypothesise that the entire island encoding *bla*_NDM-1_ was transferred from an FII pGUE-NDM-like plasmid (F2:A-:B-) to an F-type pIP1206-like plasmid (F36/F22:A1:B20) after divergence of MS6192, MS6193 and MS6194 from MS6204 (Figure 7). According to our model, the most recent common ancestor (MRCA) of MS6204, MS6192, MS6193 and MS6194 contained both an FII pGUE-NDM-like plasmid carrying the *bla*_NDM-1_ resistance island and an F-type pIP1206-like plasmid carrying a different resistance island. In the MRCA of MS6192, MS6193 and MS6194, the *bla*_NDM-1_ resistance island appears to have been transferred from the pGUE-NDM-like FII plasmid to the pIP1206-like F-type plasmid via a transposition-like event with subsequent loss of *bla*_NDM-1_ and most of the resistance island from the donor plasmid. Although we cannot formally rule out acquisition of the 21.6 Kb *bla*_NDM-1_ encoding resistance island from a different intermediary plasmid, we suggest a direct transfer between co-existing ancestral plasmids is the most parsimonious explanation for the observed plasmid sequences in this study. Whilst the ILR of Tn*5403* and IRR of Tn*As3* are present in pMS6192A-NDM, pMS6193A-NDM and pMS6194A-NDM, along with intact Tn*As3* transposase and recombinase genes, it remains unknown whether the MDR island in its current form is capable of active transposition.

**Figure 7.**
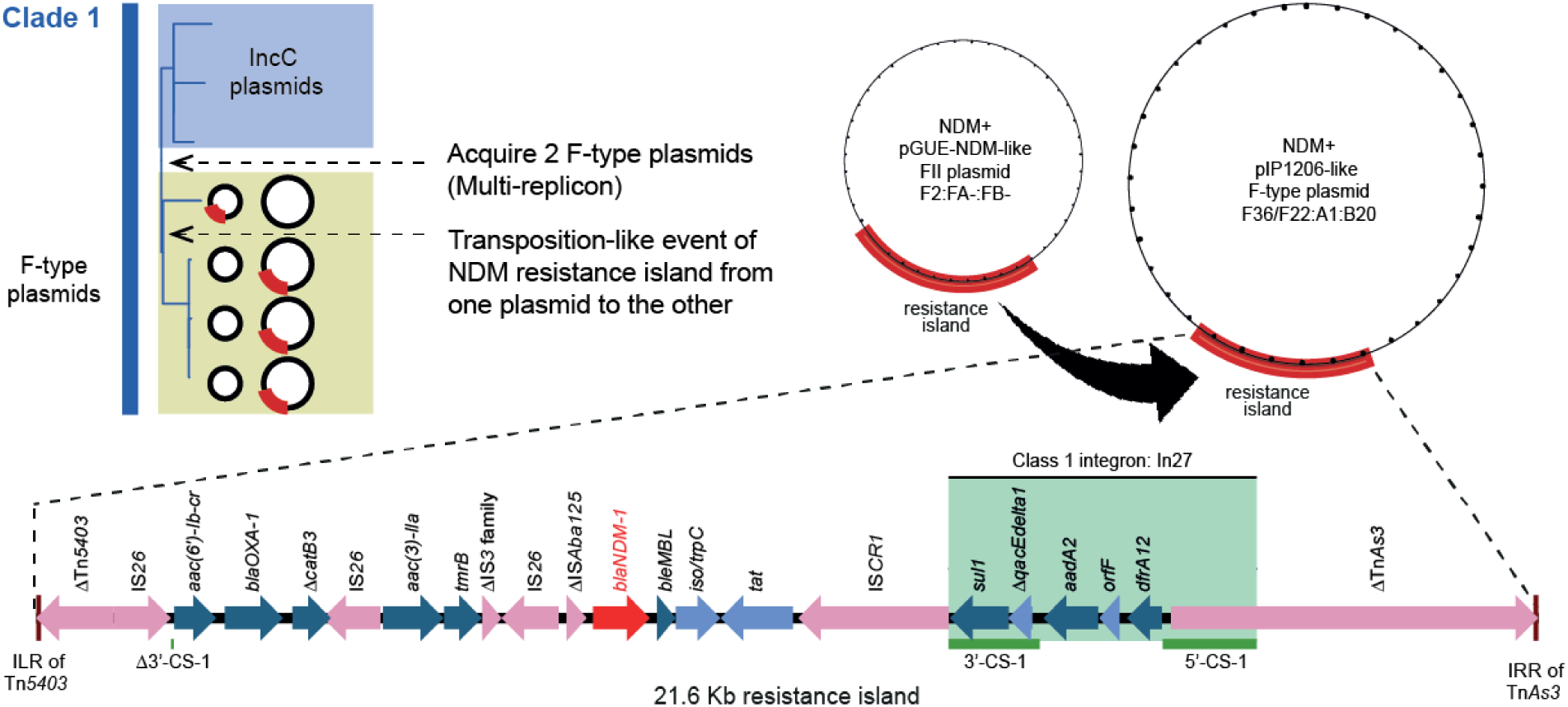
Proposed model of transfer of NDM+ resistance island between F-type plasmids. We predict that acquisition of a pGUE-NDM-like plasmid and a pIP1206-like plasmid occurred in the MRCA of MS6204, MS6192, MS6194 and MS6193. A transposition-like event of the entire resistance island between both FII plasmids resulted in the transfer of *bla*NDM-1 and surrounding resistance genes. Key regions of the resistance island are indicated: Δ: truncated gene, CDS: light blue, IS/Tns: light pink, AMR genes: teal, Integron: green rectangle. 5’ and 3-conserved sequences of class 1 integrons: dark green. The *bla*_NDM-1_ gene is labelled in red. The ILR of Tn*5403* and IRR of Tn*As3* are indicated at the edges of the island.

We note that the relatively small number of ST101 genomes used here may constrain our ability to fully characterise the clade structure and thus verify our plasmid model. However, both clades were supported in a larger analysis of 263 available ST101 draft genomes obtained from Enterobase (30/07/18) (40) that were of suitable quality for phylogenetic analysis (Supplementary Appendix, Fig S5). Additionally, our finding that *bla*_NDM-1_ (or its variants) was confined to a single sub-lineage within Clade 1A remains intact. The conservation of the pGUE-NDM-like (FII) and pIP1206-like (F-type) plasmid backbones in this larger ST101 dataset were also congruent with a single acquisition into a subclade of Clade 1A (with loss of the FII plasmid in some strains) (Supplementary Appendix, Fig S6 and Fig S7). Complete conservation of the *bla*_NDM-1_ resistance island was restricted to only a few strains, with the clade structure comparable to that of our smaller dataset, further supporting our model of island transfer between plasmids (Supplementary Appendix, Fig S8).

## Discussion

Here we used PacBio SMRT sequencing to provide the most comprehensive genomic snapshot of *bla*_NDM-1_ carriage in the *E. coli* ST101 lineage to date. We show that this lineage is monophyletic within the B1 phylogroup with at least two distinct clades that we have labelled Clade 1 and Clade 2. We show that *bla*_NDM-1_ carriage is confined to Clade 1 and associated with fluoroquinolone resistance mutations and other resistance genes such *bla*_CTX-M-15_. By determining the complete sequence of two closely related ST101 strains and the complete plasmid sequences for five other ST101 strains, we have characterised the genomic context of *bla*_NDM-1_ and other resistance genes. Notably, we revealed a mosaic region of AMR genes including *bla*_NDM-1_ within the ARI-A region of IncC plasmids and transfer of a large MDR resistance region encoding *bla*_NDM-1_ between two different F-type plasmids. These results highlight the power of long-read sequencing in revealing the full complexities of mobile genetic elements.

Identifying the genomic characteristics of *E. coli* lineages is crucial to explaining the evolution and global dissemination of successful clones. Carbapenem resistance resulting from the acquisition of plasmids carrying *bla*_NDM-1_ has been characterised in several *E. coli* clonal lineages, such as ST405, ST131 and ST101 (12). However, despite numerous reports of *bla*_NDM-1_ or variants within the *E. coli* ST101 lineage (4, 12-14, 17, 36), there remains a paucity of whole genome studies. The use of PacBio SMRT sequencing long-read technology in this study enabled the *de novo* assembly of two complete ST101 genomes (MS6192 and MS6193) and five high-quality draft genomes (MS6194, MS6201, MS6203, MS6204 and MS6207). The inability to completely resolve the genomes of the other five isolates was due to low data output achieved using a single SMRT cell on the PacBio RSI platform. Despite this impediment we could completely resolve all plasmid sequences carrying *bla*_NDM-1_ and define the context of AMR elements within these seven isolates.

Our comparisons of the very closely related ST101 strains MS6192, MS6193 and MS6194 revealed a pattern of genome conservation consistent with a close evolutionary relationship. For example, a conserved core and accessory genome with only 12 core SNP differences between the chromosomes of MS6192 and MS6193, the conservation of both FII plasmid backbones (pGUE-NDM-like and pIP1206-like), as well as the stepwise acquisition of Phi 8 (present in MS6194 and MS6193, but not MS6192) and the Tn*7*-like transposon (present only in MS6193) are consistent with epidemiologically-linked strains. This level of chromosomal similarity is consistent with long-term gut colonisation by a clonal population of Enterobacteriaceae (41). In fact, we also recently reported the existence of an indigenous clonal lineage (with significant plasmid diversity) amongst *E. coli* ST131 isolates collected from urine and fecal samples over a five-year period from a patient with recurrent urinary tract infection (42). Analysis of the core genome of the 20 ST101 isolates also indicates a close relationship, however considerable diversity is observed in their MGE complement. These differences are particularly evident between clades, where MGEs such as prophages and genomic islands defined in our reference ST101 genomes MS6192 and MS6193, in general, are not conserved. Exceptions include the GI-*leuX* locus, which is intact in most Clade 1 and 2 strains. Nonetheless, our study is limited by the availability of temporal and geographic metadata for all sequenced strains, which restricted our ability to interpret the ST101 phylogeny.

MDR is primarily disseminated within ST101 Clade 1A by plasmids of the groups Incl1, IncC and F-type. The genetic elements harbouring *bla*_NDM-1_ can spread among Gram-negative bacteria such as *Acinetobacter, Pseudomonas and Enterobacteriaceae* including *K. pneumoniae* and *E. coli* (43), where IncC and FII plasmids have previously been reported as carriers of *bla*_NDM-1_ (8, 44). Antimicrobial susceptibility testing showed that the AMR phenotype closely matched the AMR genotype for the seven PacBio sequenced ST101 genomes in this study. However, MS6207 had an intermediate resistance phenotype to meropenem and imipenem (Supplementary Dataset 1, Table S9), despite the presence of the *bla*_NDM-1_ gene, reflecting the fact that AMR prediction from WGS data in Enterobacteriaceae remains difficult (45, 46).

In this study, we characterised two pathways for *bla*_NDM-1_ persistence in ST101 Clade 1A. For strains with IncC plasmids, carriage of *bla*_NDM-1_ was associated with recombination of class 1 integrons and transposition of composite transposons and IS elements, resulting in differences in AMR gene repertoire. In contrast, we show carriage of the *bla*_NDM-1_ resistance island in an FII plasmid, with predicted transfer to a co-existing F-type plasmid. Transposition events mediated by both IS and transposons enable the mobility of entire resistance islands, particularly in the *Enterobacteriaceae* (47). For example, a recent study showed that transposition mediated by IS26, Tn*5403* and Tn*3*-family transposons such as those found flanking this *bla*_NDM-1_ resistance island facilitated plasmid rearrangements in carbapenem resistant *K. pneumoniae* (39). Although we cannot be certain of the precise events leading to the evolution of pMS6192A-NDM, pMS6193A-NDM and pMS6194A-NDM, our proposed model suggests that pGUE-NDM-like plasmids may have played significant role in early dissemination of *bla*_NDM-1_ in *E. coli*. However, we note that a lack of ST101 complete genomes in both major clades has constrained our ability to comprehensively confirm our proposed model of *bla*_NDM-1_ resistance island transfer.

Fluoroquinolone resistance in Clade 1 strains is attributed to vertically transmitted point mutations within the QRDR of *gyrA* (S83L, D87N and D87Y alleles) and *parC* (S80I allele). This is similar to the ST131 lineage where acquisition of QRDR alleles imparting fluoroquinolone resistance occurred in the MRCA prior to the divergence of Clade C (10). With the presence of chromosomally acquired fluoroquinolone resistance and the acquisition of plasmid-encoded resistance genes to more than nine different antimicrobial classes, these extensively MDR Clade 1 strains leave few choices for treatment. Older drugs such as colistin are now being used as last-resort treatments for carbapenem-resistant infections (48). In 2016 however, an ST101 strain (ICBEC72H, included in this study) containing the *mcr-1* colistin resistance gene on an IncX plasmid backbone was identified (49). With recent reports of both carbapenem and colistin resistant *E. coli* strains appearing (50, 51), we face the very real possibility of pan-drug resistant *E. coli*. Overall, these high-quality *E. coli* ST101 genomes will provide an important reference for further analysis of how mobile genetic elements and antimicrobial resistance influence the evolution of this emerging, MDR *E. coli* lineage.

## Methods

### Bacterial genome sequencing and assembly of our seven PacBio sequenced strains

The NDM-positive *E. coli* strains used in this study have been described previously and were screened for the presence of *bla*_NDM-1_ by PCR (20, 21). Additional details regarding strain selection and DNA isolation methods can be found in the Supplementary Appendix. Six of seven strains were sequenced on the PacBio RSI instrument, as previously described (24), using a 10 Kb insert library. *E. coli* ST101 strain MS6192 used six SMRT cells, with MS6194, MS6201, MS6203, MS6204, MS6207 using two SMRT cells each. *E. coli* ST101 strain MS6193 was sequenced on the PacBio RSII instrument using three SMRT cells with P4-C2 chemistry. Raw sequencing reads were *de novo* assembled using the Hierarchical Genome Assembly Process (HGAP) (52) as implemented in the PacBio SMRT Analysis software suite v2.3.0, with a seed read cut-off length of 6,000 bp and default parameters. For *E. coli* MS6192 and MS6193, contig order and orientation was determined using Contiguity and nucleotide *bla*st of the overlapping sequences at each contig end. The merged assemblies were then checked against the closely related commensal strain *E. coli* SE11 using Mauve v2.3.1 (53). Additionally, two small, spurious contigs were generated during the *de novo* assembly of MS6192, consisting of rRNA and tRNA genes that mapped to multiple locations with 100% nucleotide identity to the MS6192 chromosome. These were deemed chimeric contigs and were not included in the final assembly. Contiguity (54) was used to visualise the assembly, with overlapping contigs manually trimmed and circularised. Misassemblies were corrected by aligning the reads using *BLA*SR (55), prior to sequence polishing. Raw sequence reads were then mapped to these consensus contigs using Quiver, implemented in the SMRT Analysis software suite, filtering errors from the assembly. Furthermore, read pileups across all repetitive (rRNA) regions were manually inspected to ensure that their position was supported by spanning reads.

Draft PacBio sequenced ST101 assemblies were screened for plasmid sequences using all-versus-all nucleotide comparisons to generate inter and intra-contig pairwise alignments. All contigs were then screened for overlapping sequences at the 5’ and 3’ ends as this is an artefact of the HGAP assembly process (52) and contigs with self-similar ends likely represent fully complete, circular plasmids. One end of the contig was trimmed to produce a circular sequence. Additional, putative plasmid contigs were screened by nucleotide *BLA*ST comparison against the NCBI non-redundant database. As an additional quality control step, Illumina HiSeq paired-end reads were mapped against each respective genome using bwa 0.7.17-r1188 (56), with Pilon v1.22 (57) used to correct for small indels. Plasmid misassemblies in MS6192 were corrected using alternative assembly methods (Supplementary Appendix).

### Genome annotation

The automated annotation of all PacBio sequenced ST101 strains (2 complete, 5 draft assemblies) was done using Prokka v1.11 (58) using a custom Escherichia database consisting of protein sequences from the EcoCyc website (http://ecocyc.org/) and annotated NDM+ reference plasmids (Supplementary Dataset, Table S11). For the complete genomes: *E. coli* MS6192 and MS6193, prophages, genomic islands and insertion sequences (IS) were identified using PHAST (59), IslandViewer 3 (60) and ISSaga (61) respectively. Draft annotations for MS6192 and MS6193 were then visualised using Artemis (62) and subjected to manual improvement. Insertion sequence, integrons and antimicrobial resistance cassettes within the complete plasmid sequences were manually curated by nucleotide comparisons against ISSaga, Integrall and the Repository of Antibiotic Resistance Cassettes (61, 63, 64) databases, respectively.

### Additional *E. coli* ST101 genomes included in this study

To further characterise the ST101 lineage, 13 additional ST101 draft genomes were downloaded from GenBank or the SRA. Three genomes (ECO0117, eo1667 and PI7) were only available as Illumina paired-end short reads and were assembled *de-novo* using Spades v3.10.1 with default parameters. Strain names, accessions, sources, sequencing method and available metadata are summarised in Supplementary Dataset, Table S3.

### Phylogenetic analysis and recombination filtering

To determine the phylogenetic relationship of ST101 and other *E. coli* lineages, core-genome single nucleotide polymorphism (SNP) trees were constructed. A core-genome SNP alignment of the 20 ST101 strains and an additional 65 representative complete *E. coli* genomes (Supplementary Dataset, Table S12) was produced by parSNP v1.2, implemented in the Harvest suite (65) and aligned against the well-characterised ExPEC complete genome *E. coli* EC958 reference (23, 24) defining 182,264 core SNPs. To integrate the ST101 draft and complete genomes into our phylogenomic analyses and account for differences between short and long read technologies, error-free 71-bp paired-end Illumina reads were simulated from the draft assemblies and complete genomes and used as input to produce a whole genome SNP alignment using Snippy v4.3.5 (https://github.com/tseemann/snippy.git) against the ST101 complete genome MS6192. A pseudogenome of this ST101 SNP alignment was used as input for Gubbins v2.3.1 (66), which detects recombination in closely related groups of isolates. Gubbins was run using default settings in “raxml mode” generating a phylogenetic tree with a generalized time-reversible model and gamma correction (GTRGAMMA). This generated a recombination-filtered core-SNP alignment of 2,106 SNPs. Maximum likelihood (ML) phylogenetic trees were then constructed from both core-genome SNP-based alignments with RAxML 8.1.15 (67) using the GTRGAMMA model and 1000 bootstrap replicates. The resulting ML phylogenetic trees were visualised using FigTree v1.4.2 (http://tree.bio.ed.ac.uk/software/figtree/).

### Comparative genomics of *E. coli* ST101

For all *E. coli* ST101 genomes included in this study, acquired AMR genes were functionally annotated by nucleotide comparisons (a cutoff of 90% identity over >=60% query cover) against ResFinder v2.1 (68). O antigen and H antigen typing were determined using SeroType Finder v1.1 (69) and the EcOH database (70) within SRST2 (71), with a nucleotide ID threshold of 85%. Additionally, sequences were screened for the presence of mutations in the QRDR of gyrA and parC. For each isolate and the *E. coli* K12 strain MG1655 reference genome (Genbank accession: U00096), the amino acid sequences of GyrA and ParC were aligned in MEGA v6.06 (72), using MUSCLE (73) and default parameters. All isolates also underwent *fimH in silico* typing using FimTyper v1.0 (https://cge.cbs.dtu.dk/services/FimTyper/). For plasmid analyses, the plasmid replicons were detected using PlasmidFinder v1.3 with a nucleotide ID threshold of 90% and the plasmid Multi-Locus Sequence Typing (pMLST) (*Enterobacteriaceae*) was determined using pMLST v1.4 (74). IncC pMLST was determined *in silico* against the four essential IncC genes *repA, parA, parB* and *053* (34). BRIG v.0.95 (75), EasyFig v2.2.2 (76) and Phandango v1.1.0 (77) were used to visually compare the genomes. Methods for conjugation assays, phenotypic antimicrobial susceptibility testing and plasmid mating assays can be found in the Supplementary Appendix.

### Data

Complete genomes of MS6192 and MS6193, draft genome assemblies and PacBio and Illumina sequence read data for MS6192, MS6193, MS6194, MS6201, MS6203, MS6204 and MS6207 are available under the BioProjects: PRJNA580334, PRJNA580336, PRJNA580337, PRJNA580338, PRJNA580339, PRJNA580341 and PRJNA580340 respectively.

## Acknowledgements

Author contributions: Conceptualisation: MMA, BMF, MAS and SAB. Investigation: MMA and BMF. Formal Analysis: MMA. MDP, KMP, AMH, SJH and DLP assisted in clinical/wet-lab experiments. BMF, SJH, LWR and RTW assisted in data analysis. KGC, TMC and WFY performed the PacBio sequencing. Illumina sequencing was performed at the Australian Genome Research Facility (AGRF), The University of Queensland, Australia. Resources: TRW, KGC, MAS and SAB. Supervision: BMF, MAS and SAB. Writing (Original Draft Preparation): MMA, BMF and SAB. Writing (Review and Editing): MMA, MDP, AMH, SJH, KGC, MAS and SAB. All authors contributed to the final review and edits.

## References

1. Poolman JT, Wacker M. 2016. Extraintestinal Pathogenic Escherichia coli, a Common Human Pathogen: Challenges for Vaccine Development and Progress in the Field. J Infect Dis 213:6–13.

2. Yong D, Toleman MA, Giske CG, Cho HS, Sundman K, Lee K, Walsh TR. 2009. Characterization of a new metallo-beta-lactamase gene, bla(NDM-1), and a novel erythromycin esterase gene carried on a unique genetic structure in Klebsiella pneumoniae sequence type 14 from India. Antimicrob Agents Chemother 53:5046–54.

3. Nordmann P, Poirel L, Walsh TR, Livermore DM. 2011. The emerging NDM carbapenemases. Trends Microbiol 19:588–95.

4. Peirano G, Mulvey GL, Armstrong GD, Pitout JD. 2013. Virulence potential and adherence properties of Escherichia coli that produce CTX-M and NDM beta-lactamases. J Med Microbiol 62:525–30.

5. Woodford N, Turton JF, Livermore DM. 2011. Multiresistant Gram-negative bacteria: the role of high-risk clones in the dissemination of antibiotic resistance. FEMS Microbiol Rev 35:736–55.

6. Zarrilli R, Pournaras S, Giannouli M, Tsakris A. 2013. Global evolution of multidrug-resistant Acinetobacter baumannii clonal lineages. Int J Antimicrob Agents 41:11–9.

7. Coque TM, Novais A, Carattoli A, Poirel L, Pitout J, Peixe L, Baquero F, Canton R, Nordmann P. 2008. Dissemination of clonally related Escherichia coli strains expressing extended-spectrum beta-lactamase CTX-M-15. Emerg Infect Dis 14:195–200.

8. Bonnin RA, Poirel L, Carattoli A, Nordmann P. 2012. Characterization of an IncFII plasmid encoding NDM-1 from Escherichia coli ST131. PLoS One 7:e34752.

9. Petty NK, Ben Zakour NL, Stanton-Cook M, Skippington E, Totsika M, Forde BM, Phan MD, Gomes Moriel D, Peters KM, Davies M, Rogers BA, Dougan G, Rodriguez-Bano J, Pascual A, Pitout JD, Upton M, Paterson DL, Walsh TR, Schembri MA, Beatson SA. 2014. Global dissemination of a multidrug resistant Escherichia coli clone. Proc Natl Acad Sci U S A 111:5694–9.

10. Ben Zakour NL, Alsheikh-Hussain AS, Ashcroft MM, Khanh Nhu NT, Roberts LW, Stanton-Cook M, Schembri MA, Beatson SA. 2016. Sequential Acquisition of Virulence and Fluoroquinolone Resistance Has Shaped the Evolution of Escherichia coli ST131. MBio 7:e00347–16.

11. Price LB, Johnson JR, Aziz M, Clabots C, Johnston B, Tchesnokova V, Nordstrom L, Billig M, Chattopadhyay S, Stegger M, Andersen PS, Pearson T, Riddell K, Rogers P, Scholes D, Kahl B, Keim P, Sokurenko EV. 2013. The epidemic of extended-spectrum-beta-lactamase-producing Escherichia coli ST131 is driven by a single highly pathogenic subclone, H30-Rx. MBio 4:e00377–13.

12. Mushtaq S, Irfan S, Sarma JB, Doumith M, Pike R, Pitout J, Livermore DM, Woodford N. 2011. Phylogenetic diversity of Escherichia coli strains producing NDM-type carbapenemases. J Antimicrob Chemother 66:2002–5.

13. Yoo JS, Kim HM, Koo HS, Yang JW, Yoo JI, Kim HS, Park HK, Lee YS. 2013. Nosocomial transmission of NDM-1-producing Escherichia coli ST101 in a Korean hospital. J Antimicrob Chemother 68:2170–2.

14. Poirel L, Savov E, Nazli A, Trifonova A, Todorova I, Gergova I, Nordmann P. 2014. Outbreak caused by NDM-1- and RmtB-producing Escherichia coli in Bulgaria. Antimicrob Agents Chemother 58:2472–4.

15. Mantilla-Calderon D, Jumat MR, Wang T, Ganesan P, Al-Jassim N, Hong PY. 2016. Isolation and Characterization of NDM-Positive Escherichia coli from Municipal Wastewater in Jeddah, Saudi Arabia. Antimicrob Agents Chemother 60:5223–31.

16. Mora A, Blanco M, Lopez C, Mamani R, Blanco JE, Alonso MP, Garcia-Garrote F, Dahbi G, Herrera A, Fernandez A, Fernandez B, Agulla A, Bou G, Blanco J. 2011. Emergence of clonal groups O1:HNM-D-ST59, O15:H1-D-ST393, O20:H34/HNM-D-ST354, O25b:H4-B2-ST131 and ONT:H21,42-B1-ST101 among CTX-M-14-producing Escherichia coli clinical isolates in Galicia, northwest Spain. Int J Antimicrob Agents 37:16–21.

17. Ranjan A, Shaik S, Mondal A, Nandanwar N, Hussain A, Semmler T, Kumar N, Tiwari SK, Jadhav S, Wieler LH, Ahmed N. 2016. Molecular Epidemiology and Genome Dynamics of New Delhi Metallo-beta-Lactamase-Producing Extraintestinal Pathogenic Escherichia coli Strains from India. Antimicrob Agents Chemother 60:6795–6805.

18. Conlan S, Thomas PJ, Deming C, Park M, Lau AF, Dekker JP, Snitkin ES, Clark TA, Luong K, Song Y, Tsai YC, Boitano M, Dayal J, Brooks SY, Schmidt B, Young AC, Thomas JW, Bouffard GG, Blakesley RW, Mullikin JC, Korlach J, Henderson DK, Frank KM, Palmore TN, Segre JA. 2014. Single-molecule sequencing to track plasmid diversity of hospital-associated carbapenemase-producing Enterobacteriaceae. Sci Transl Med 6:254ra126.

19. Zowawi HM, Forde BM, Alfaresi M, Alzarouni A, Farahat Y, Chong TM, Yin WF, Chan KG, Li J, Schembri MA, Beatson SA, Paterson DL. 2015. Stepwise evolution of pandrug-resistance in Klebsiella pneumoniae. Sci Rep 5:15082.

20. Djoko KY, Achard MES, Phan MD, Lo AW, Miraula M, Prombhul S, Hancock SJ, Peters KM, Sidjabat H, Harris PN, Mitic N, Walsh TR, Anderson GJ, Shafer WM, Paterson DL, Schenk G, McEwan AG, Schembri MA. 2017. Copper ions and coordination complexes as novel carbapenem adjuvants. Antimicrob Agents Chemother doi:10.1128/aac.02280-17.

21. Kumarasamy KK, Toleman MA, Walsh TR, Bagaria J, Butt F, Balakrishnan R, Chaudhary U, Doumith M, Giske CG, Irfan S, Krishnan P, Kumar AV, Maharjan S, Mushtaq S, Noorie T, Paterson DL, Pearson A, Perry C, Pike R, Rao B, Ray U, Sarma JB, Sharma M, Sheridan E, Thirunarayan MA, Turton J, Upadhyay S, Warner M, Welfare W, Livermore DM, Woodford N. 2010. Emergence of a new antibiotic resistance mechanism in India, Pakistan, and the UK: a molecular, biological, and epidemiological study. Lancet Infect Dis 10:597–602.

22. Oshima K, Toh H, Ogura Y, Sasamoto H, Morita H, Park SH, Ooka T, Iyoda S, Taylor TD, Hayashi T, Itoh K, Hattori M. 2008. Complete genome sequence and comparative analysis of the wild-type commensal Escherichia coli strain SE11 isolated from a healthy adult. DNA Res 15:375–86.

23. Totsika M, Beatson SA, Sarkar S, Phan MD, Petty NK, Bachmann N, Szubert M, Sidjabat HE, Paterson DL, Upton M, Schembri MA. 2011. Insights into a multidrug resistant Escherichia coli pathogen of the globally disseminated ST131 lineage: genome analysis and virulence mechanisms. PLoS One 6:e26578.

24. Forde BM, Ben Zakour NL, Stanton-Cook M, Phan MD, Totsika M, Peters KM, Chan KG, Schembri MA, Upton M, Beatson SA. 2014. The complete genome sequence of Escherichia coli EC958: a high quality reference sequence for the globally disseminated multidrug resistant E. coli O25b:H4-ST131 clone. PLoS One 9:e104400.

25. Alteri CJ, Mobley HL. 2016. The Versatile Type VI Secretion System. Microbiol Spectr 4.

26. Ren CP, Beatson SA, Parkhill J, Pallen MJ. 2005. The Flag-2 locus, an ancestral gene cluster, is potentially associated with a novel flagellar system from Escherichia coli. J Bacteriol 187:1430–40.

27. Zhou M, Guo Z, Duan Q, Hardwidge PR, Zhu G. 2014. Escherichia coli type III secretion system 2: a new kind of T3SS? Vet Res 45:32.

28. Hopkins KL, Davies RH, Threlfall EJ. 2005. Mechanisms of quinolone resistance in Escherichia coli and Salmonella: recent developments. Int J Antimicrob Agents 25:358–73.

29. Johnson TJ, Elnekave E, Miller EA, Munoz-Aguayo J, Flores Figueroa C, Johnston B, Nielson DW, Logue CM, Johnson JR. 2019. Phylogenomic Analysis of Extraintestinal Pathogenic Escherichia coli Sequence Type 1193, an Emerging Multidrug-Resistant Clonal Group. Antimicrob Agents Chemother 63.

30. Dortet L, Nordmann P, Poirel L. 2012. Association of the emerging carbapenemase NDM-1 with a bleomycin resistance protein in Enterobacteriaceae and Acinetobacter baumannii. Antimicrob Agents Chemother 56:1693–7.

31. Campos JC, da Silva MJF, dos Santos PRN, Barros EM, Pereira MdO, Seco BMS, Magagnin CM, Leiroz LK, de Oliveira TGM, de Faria-Júnior C, Cerdeira LT, Barth AL, Sampaio SCF, Zavascki AP, Poirel L, Sampaio JLM. 2015. Characterization of Tn3000, a Transposon Responsible for bla(NDM-1) Dissemination among Enterobacteriaceae in Brazil, Nepal, Morocco, and India. Antimicrobial Agents and Chemotherapy 59:7387–7395.

32. Marquez-Ortiz RA, Haggerty L, Olarte N, Duarte C, Garza-Ramos U, Silva-Sanchez J, Castro BE, Sim EM, Beltran M, Moncada MV, Valderrama A, Castellanos JE, Charles IG, Vanegas N, Escobar-Perez J, Petty NK. 2017. Genomic Epidemiology of NDM-1-Encoding Plasmids in Latin American Clinical Isolates Reveals Insights into the Evolution of Multidrug Resistance. Genome Biol Evol 9:1725–1741.

33. Sugawara Y, Akeda Y, Sakamoto N, Takeuchi D, Motooka D, Nakamura S, Hagiya H, Yamamoto N, Nishi I, Yoshida H, Okada K, Zin KN, Aye MM, Tonomo K, Hamada S. 2017. Genetic characterization of blaNDM-harboring plasmids in carbapenem-resistant Escherichia coli from Myanmar. PLoS One 12:e0184720.

34. Hancock SJ, Phan MD, Peters KM, Forde BM, Chong TM, Yin WF, Chan KG, Paterson DL, Walsh TR, Beatson SA, Schembri MA. 2016. Identification of IncA/C Plasmid Replication and Maintenance Genes and Development of a Plasmid Multi-Locus Sequence-Typing Scheme. Antimicrob Agents Chemother doi:10.1128/aac.01740-16.

35. Fiett J, Baraniak A, Izdebski R, Sitkiewicz I, Zabicka D, Meler A, Filczak K, Hryniewicz W, Gniadkowski M. 2014. The first NDM metallo-beta-lactamase-producing Enterobacteriaceae isolate in Poland: evolution of IncFII-type plasmids carrying the bla(NDM-1) gene. Antimicrob Agents Chemother 58:1203–7.

36. Toleman MA, Bugert JJ, Nizam SA. 2015. Extensively drug-resistant New Delhi metallo-beta-lactamase-encoding bacteria in the environment, Dhaka, Bangladesh, 2012. Emerg Infect Dis 21:1027–30.

37. Wailan AM, Paterson DL, Kennedy K, Ingram PR, Bursle E, Sidjabat HE. 2015. Genomic Characteristics of NDM-Producing Enterobacteriaceae Isolates in Australia and Their blaNDM Genetic Contexts. Antimicrob Agents Chemother 60:136–41.

38. Siguier P, Perochon J, Lestrade L, Mahillon J, Chandler M. 2006. ISfinder: the reference centre for bacterial insertion sequences. Nucleic Acids Res 34:D32–6.

39. He S, Chandler M, Varani AM, Hickman AB, Dekker JP, Dyda F. 2016. Mechanisms of Evolution in High-Consequence Drug Resistance Plasmids. MBio 7.

40. Alikhan NF, Zhou Z, Sergeant MJ, Achtman M. 2018. A genomic overview of the population structure of Salmonella. PLoS Genet 14:e1007261.

41. Conlan S, Park M, Deming C, Thomas PJ, Young AC, Coleman H, Sison C, Weingarten RA, Lau AF, Dekker JP, Palmore TN, Frank KM, Segre JA. 2016. Plasmid Dynamics in KPC-Positive Klebsiella pneumoniae during Long-Term Patient Colonization. MBio 7.

42. Forde BM, Roberts LW, Phan MD, Peters KM, Fleming BA, Russell CW, Lenherr SM, Myers JB, Barker AP, Fisher MA, Chong TM, Yin WF, Chan KG, Schembri MA, Mulvey MA, Beatson SA. 2019. Population dynamics of an Escherichia coli ST131 lineage during recurrent urinary tract infection. Nat Commun 10:3643.

43. Diene SM, Rolain JM. 2014. Carbapenemase genes and genetic platforms in Gram-negative bacilli: Enterobacteriaceae, Pseudomonas and Acinetobacter species. Clin Microbiol Infect 20:831–8.

44. Hudson CM, Bent ZW, Meagher RJ, Williams KP. 2014. Resistance determinants and mobile genetic elements of an NDM-1-encoding Klebsiella pneumoniae strain. PLoS One 9:e99209.

45. Ingle DJ, Levine MM, Kotloff KL, Holt KE, Robins-Browne RM. 2018. Dynamics of antimicrobial resistance in intestinal Escherichia coli from children in community settings in South Asia and sub-Saharan Africa. Nat Microbiol 3:1063–1073.

46. Su M, Satola SW, Read TD. 2019. Genome-Based Prediction of Bacterial Antibiotic Resistance. J Clin Microbiol 57.

47. Toleman MA, Walsh TR. 2011. Combinatorial events of insertion sequences and ICE in Gram-negative bacteria. FEMS Microbiol Rev 35:912–35.

48. Yamamoto M, Pop-Vicas AE. 2014. Treatment for infections with carbapenem-resistant Enterobacteriaceae: what options do we still have? Crit Care 18:229.

49. Fernandes MR, McCulloch JA, Vianello MA, Moura Q, Perez-Chaparro PJ, Esposito F, Sartori L, Dropa M, Matte MH, Lira DP, Mamizuka EM, Lincopan N. 2016. First Report of the Globally Disseminated IncX4 Plasmid Carrying the mcr-1 Gene in a Colistin-Resistant Escherichia coli Sequence Type 101 Isolate from a Human Infection in Brazil. Antimicrob Agents Chemother 60:6415–7.

50. Zheng B, Dong H, Xu H, Lv J, Zhang J, Jiang X, Du Y, Xiao Y, Li L. 2016. Coexistence of MCR-1 and NDM-1 in Clinical Escherichia coli Isolates. Clin Infect Dis 63:1393–1395.

51. Zhong LL, Zhang YF, Doi Y, Huang X, Zhang XF, Zeng KJ, Shen C, Patil S, Xing Y, Zou Y, Tian GB. 2017. Coproduction of MCR-1 and NDM-1 by Colistin-Resistant Escherichia coli Isolated from a Healthy Individual. Antimicrob Agents Chemother 61.

52. Chin CS, Alexander DH, Marks P, Klammer AA, Drake J, Heiner C, Clum A, Copeland A, Huddleston J, Eichler EE, Turner SW, Korlach J. 2013. Nonhybrid, finished microbial genome assemblies from long-read SMRT sequencing data. Nat Methods 10:563–9.

53. Darling AE, Mau B, Perna NT. 2010. progressiveMauve: multiple genome alignment with gene gain, loss and rearrangement. PLoS One 5:e11147.

54. Sullivan MJ, L. Bzn, M. FB, Stanton-Cook M, Beatson SA. 2015. Contiguity: Contig adjacency graph construction and visualisation. PeerJ PrePrints 3:e1273.

55. Chaisson MJ, Tesler G. 2012. Mapping single molecule sequencing reads using basic local alignment with successive refinement (BLASR): application and theory. BMC Bioinformatics 13:238.

56. Li H, Durbin R. 2009. Fast and accurate short read alignment with Burrows-Wheeler transform. Bioinformatics 25:1754–60.

57. Walker BJ, Abeel T, Shea T, Priest M, Abouelliel A, Sakthikumar S, Cuomo CA, Zeng Q, Wortman J, Young SK, Earl AM. 2014. Pilon: An Integrated Tool for Comprehensive Microbial Variant Detection and Genome Assembly Improvement. PLOS ONE 9:e112963.

58. Seemann T. 2014. Prokka: rapid prokaryotic genome annotation. Bioinformatics 30:2068-9. 59.

59. Zhou Y, Liang Y, Lynch KH, Dennis JJ, Wishart DS. 2011. PHAST: a fast phage search tool. Nucleic Acids Res 39:W347–52.

60. Dhillon BK, Laird MR, Shay JA, Winsor GL, Lo R, Nizam F, Pereira SK, Waglechner N, McArthur AG, Langille MG, Brinkman FS. 2015. IslandViewer 3: more flexible, interactive genomic island discovery, visualization and analysis. Nucleic Acids Res 43:W104–8.

61. Varani AM, Siguier P, Gourbeyre E, Charneau V, Chandler M. 2011. ISsaga is an ensemble of web-based methods for high throughput identification and semi-automatic annotation of insertion sequences in prokaryotic genomes. Genome Biol 12:R30.

62. Carver T, Harris SR, Berriman M, Parkhill J, McQuillan JA. 2012. Artemis: an integrated platform for visualization and analysis of high-throughput sequence-based experimental data. Bioinformatics 28:464–9.

63. Moura A, Soares M, Pereira C, Leitao N, Henriques I, Correia A. 2009. INTEGRALL: a database and search engine for integrons, integrases and gene cassettes. Bioinformatics 25:1096–8.

64. Tsafnat G, Copty J, Partridge SR. 2011. RAC: Repository of Antibiotic resistance Cassettes. Database (Oxford) 2011:bar054.

65. Treangen TJ, Ondov BD, Koren S, Phillippy AM. 2014. The Harvest suite for rapid core-genome alignment and visualization of thousands of intraspecific microbial genomes. Genome Biol 15:524.

66. Croucher NJ, Page AJ, Connor TR, Delaney AJ, Keane JA, Bentley SD, Parkhill J, Harris SR. 2015. Rapid phylogenetic analysis of large samples of recombinant bacterial whole genome sequences using Gubbins. Nucleic Acids Res 43:e15.

67. Stamatakis A. 2006. RAxML-VI-HPC: maximum likelihood-based phylogenetic analyses with thousands of taxa and mixed models. Bioinformatics 22:2688–90.

68. Zankari E, Hasman H, Cosentino S, Vestergaard M, Rasmussen S, Lund O, Aarestrup FM, Larsen MV. 2012. Identification of acquired antimicrobial resistance genes. J Antimicrob Chemother 67:2640–4.

69. Joensen KG, Tetzschner AM, Iguchi A, Aarestrup FM, Scheutz F. 2015. Rapid and Easy In Silico Serotyping of Escherichia coli Isolates by Use of Whole-Genome Sequencing Data. J Clin Microbiol 53:2410–26.

70. Ingle DJ, Valcanis M, Kuzevski A, Tauschek M, Inouye M, Stinear T, Levine MM, Robins-Browne RM, Holt KE. 2016. In silico serotyping of E. coli from short read data identifies limited novel O-loci but extensive diversity of O: H serotype combinations within and between pathogenic lineages. Microbial Genomics 2.

71. Inouye M, Dashnow H, Raven LA, Schultz MB, Pope BJ, Tomita T, Zobel J, Holt KE. 2014. SRST2: Rapid genomic surveillance for public health and hospital microbiology labs. Genome Med 6:90.

72. Tamura K, Stecher G, Peterson D, Filipski A, Kumar S. 2013. MEGA6: Molecular Evolutionary Genetics Analysis version 6.0. Mol Biol Evol 30:2725–9.

73. Edgar RC. 2004. MUSCLE: a multiple sequence alignment method with reduced time and space complexity. BMC Bioinformatics 5:113.

74. Carattoli A, Zankari E, Garcia-Fernandez A, Voldby Larsen M, Lund O, Villa L, Moller Aarestrup F, Hasman H. 2014. In silico detection and typing of plasmids using PlasmidFinder and plasmid multilocus sequence typing. Antimicrob Agents Chemother 58:3895–903.

75. Alikhan NF, Petty NK, Ben Zakour NL, Beatson SA. 2011. BLAST Ring Image Generator (BRIG): simple prokaryote genome comparisons. BMC Genomics 12:402.

76. Sullivan MJ, Petty NK, Beatson SA. 2011. Easyfig: a genome comparison visualizer. Bioinformatics 27:1009–10.

77. Hadfield J, Croucher NJ, Goater RJ, Abudahab K, Aanensen DM, Harris SR. 2018. Phandango: an interactive viewer for bacterial population genomics. Bioinformatics 34:292–293.

