## Supplementary Appendix for "Genomic characterisation and context of the *bla*_NDM-1_ carbapenemase in *Escherichia coli* ST101"

Supplementary Appendix Contents

Supplementary Methods	3

Strain characterisation and genomic DNA isolation.	3

Antimicrobial Susceptibility Testing of the seven PacBio sequenced ST101 strains.	3

Conjugation and Plasmid mating assay.	4

Correcting plasmid misassemblies.	5

Supplementary Results	6

*E. coli* ST101 is associated with the carriage of the *bla*_NDM-1_ gene.	6

The *bla*_CTX-M-15_ gene has been acquired and mobilised by IS*Ecp1* within Clade 1.	7

The seven PacBio sequenced NDM-positive ST101 strains are multidrug resistant.	7

Deletions in the IncC plasmid backbone alter conjugation efficiency in two *E. coli* ST101 strains.	8

IncC ARI-A regions are highly variable within IncC plasmids pMS6201A-NDM, pMS6203-NDM and pMS6207-NDM.	9

Supplementary Figures.	10

Fig S1. Whole genome nucleotide pairwise comparisons between *E. coli* ST101 strains MS6192 and MS6193.	10

Fig S2. Comparisons of *E. coli* ST101 Clade 1 IncC plasmids with pNDM-US-2.	11

Fig S3. Comparisons of NDM-1-positive F-type plasmids from *E. coli* ST101 Clade 1 strains.	12

Fig S4. Comparisons of plasmid types with a focus on the F2:A-:B- pGUE-NDM backbone and the *bla*_NDM-1_ resistance island.	13

Fig S5. Maximum Likelihood phylogenetic relationship of 283 *E. coli* ST101 genomes.	14

Fig S6. Conservation of the pGUE-NDM-like FII (F2:A-:B-) plasmid backbone in 283 *E. coli* ST101 genomes.	15

Fig S7. Conservation of the pIP1206-like F-type (F36/F22:A1:B20) plasmid backbone in 283 *E. coli* ST101 genomes.	16

Fig S8. Conservation of the *bla*_NDM-1_ resistance island in 283 *E. coli* ST101 genomes.	17

References	18

### Supplementary Methods

#### Strain characterisation and genomic DNA isolation.

The *bla*_NDM-1_-positive *E. coli* strains used in this study have been described previously (1, 2). Strains were cultured overnight in LB broth at 37**°**C for DNA isolation. Genomic DNA (gDNA) was extracted from overnight cultures using the Qiagen DNeasy Blood and Tissue kit as per the manufacturer’s instructions. This high-quality DNA was used for PCR amplification and whole genome sequencing (see below). The phylogenetic PCR screen was conducted on *chuA*, *yjaA* and *TSPE4.C2* as previously described (3). Multilocus sequence typing of *E. coli* isolates was done as per the Environmental Research Institute (ERI), University College Cork (UCC), Ireland (<http://mlst.ucc.ie/mlst/dbs/Ecoli/documents/primersColi_html>) (4). Briefly, seven housekeeping genes (*adk*, *fumC*, *gyrB*, *icd*, *mdh*, *purA* and *recA*) of the MLST scheme were amplified by PCR using Taq DNA polymerase (NEB) and sequenced by BigDye Terminator kit v3.1 (Applied Biosystems, CA, USA). The sequence types were assigned based on the MLST database from ERI, UCC (<https://pubmlst.org/escherichia/>). The virulence PCR screen was conducted on 16 genes as previously described (5) (refer Supplementary Dataset, Table S1 for exact gene designations). The presence of *bla*_NDM-1_ (sequence as per (6)) was confirmed using a PCR screen of a 621 bp segment with the following primers: 3073-NDM-F: 5’‑GGTTTGGCGATCTGGTTTTC-3’ and 3074-NDM-R: 5’-CGGAATGGCTCATCACGATC-3’. Isolates were then selected for whole genome sequencing based on their *E. coli* B1-ST101 designation and the presence of *bla*_NDM-1_.

#### Antimicrobial Susceptibility Testing of the seven PacBio sequenced ST101 strains.

Broth microdilution was performed using custom-made Sensititre plates sourced from Thermo Fisher. In brief, isolates were subbed from beads kept at -80 °C to Horse Blood Agar (HBA, BioMerieux) and incubated at 37 °C O_2_ for 24 hours. After assessing for purity, single colonies were inoculated into sterile Mueller Hinton Broth (MHB, Thermo Fisher) to produce a final concentration of 5 x 10^5^ colony forming units/ml. 50 µL was added to each well, with at least one well on each plate acting as a growth control and not containing any antibiotics. Purity plates and colony counts were performed at the time of inoculation of the 96-well plates from the MHB inoculum. The purity and count plates, as well as the 96-well sensititre plates were sealed with plastic film and incubated at 37 °C O_2_ for 24 hours. Plates were read using the Sensititre manual viewer, with growth recorded as turbidity or as a deposit of cells at the bottom of the well. The MIC was interpreted as the lowest concentration of an antimicrobial that inhibits visual growth. For T	rimethoprim-Sulfamethoxazole, the end point of growth was recorded as the well in which there was an 80% reduction in growth compared to the growth control well. Only plates with a positive growth control and that passed quality control for purity and plate counts were recorded.

#### Conjugation and Plasmid mating assay.

Conjugation assays were performed as previously described (7). For all plasmid mating assays, the sodium azide resistant *E. coli* J53 strain (8, 9) was used as recipient. *E. coli* ST101 strains MS6201, MS6203 and MS6207 were used as donor strains. Early stationary phase cultures were mixed at a ratio of 1:10 donors to recipients in 500µl LB and incubated at 37°C for 2 hours. Total colony forming units (CFUs) of donors, recipients and transconjugants were enumerated on LB agar plates with meropenem (1µg/ml), sodium azide (100µg/ml) and meropenem/sodium azide, respectively. Conjugation frequency was calculated as ‘number of transconjugants per donor’. Antibiotic resistance profiling and minimum inhibitory concentration (MIC) testing was performed as per Clinical and Laboratory Standards Insitute (CLSI) guidelines (<https://clsi.org/standards/products/microbiology/documents/m100/>). Each strain was assayed in triplicate.

#### Correcting plasmid misassemblies.

Mapping the corrected PacBio reads (via Minimap2 (10)) and the Illumina raw reads (via bwa mem v0.7.17-r1188 (11)) to the MS6192 genome assembly, identified regions of high SNP density within the plasmids compared to the consensus sequence. Region 1 (pMS6192A-NDM: positions 7,500 to 10,800 bp) showed homology with regions in both pMS6192B and pMS6192C, with region 2 identified in pMS6192C between positions 31,500 to 37,400 bp. In contrast, MS6193 (which lacks a plasmid homologous to pMS6192C and was sequenced on the RSII), did not suffer from the same suspected errors. In an attempt to resolve these misassemblies we generated a multi-fasta file of all three plasmid sequences (pMS6192A-NDM, pMS6192B and pMS6192C) and subsequently mapped the corrected PacBio reads using Minimap2 with default parameters. For misassembly region 1, we extracted all reads that mapped to this region plus 5Kb flanking the misassembly using Samtools v0.1.18 (12). The reads were then converted from bam format to fastq format using the -bamtofastq flag of Bedtools v2.292.2 (13). We next re-assembled this region using just the reads that mapped to this misassembly region with Flye v2.7.1 (14), with 5 rounds of polishing and default parameters. This newly assembled contig was then merged into the original pMS6192A-NDM assembly, replacing the misassembled region. The corrected PacBio reads were then re-mapped to this new pMS6192A-NDM assembly using Minimap2 and visualised in Artemis (15). No further SNP clusters could be observed.

Similarly, for misassembly region 2, we extracted all reads that mapped to this region, plus 5Kb flanking the misassembly and reassembled this region using just the reads that mapped to this misassembly region with Flye. However, after re-mapping the corrected PacBio reads to the new pMS6192C assembly, the high-density SNP cluster was not resolved. Further examination of this region identified numerous clipped reads, suggesting incorrect mapping of these reads. Thus, clipped reads were filtered using Samtools, with this new set of reads used to generate a new assembly using Flye. Mapping of the filtered reads to the newly generated assembly with Minimap2 identified no further SNP clusters, thus this newly assembled contig was integrated into the original pMS6192C assembly.

Comparisons of pMS6193A-NDM and pMS6194A-NDM showed that pMS6194A-NDM was 735 bp larger, therefore the raw Illumina reads of MS6194 were mapped to the pMS6194A-NDM assembly and visualised in Artemis. This showed the absence of reads in a 344 bp region: 13,200..13,541, thus this sequence was manually deleted. However, the pMS6194A-NDM assembly was still 391 bp larger than pMS6193A-NDM, so we also mapped the MS6194 raw PacBio reads to the assembly using Minimap2. Visualisation of these mapped reads using Artemis showed clipping of reads at position 13,199 bp. Further, comparisons between pMS6193A-NDM and pMS6194A-NDM using the Artemis Comparison Tool (16) showed a clear 391 bp duplication upstream of MS6194_A00020. This sequence was manually deleted and the updated pMS6194A-NDM assembly was verified by again mapping all MS6194 PacBio raw reads using Minimap2 and visualised in Artemis.

### Supplementary Results

#### *E. coli* ST101 is associated with the carriage of the *bla*_NDM-1_ gene.

We examined the sequence type (ST) and phylogeny of 16 *bla*_NDM-1_-positive *E. coli* strains isolated from India and the United Kingdom (1, 2). Five STs were identified, of which ST101 was the most predominant: ST101 (n=7; 43.8%), ST648 (n=4; 25%), ST405 (n=2; 12.5%), ST410 (n=2; 12.5%) and ST1196 (n=1; 6.2%) (Supplementary Dataset, Table S1). Phylogenetic typing by PCR was also performed and revealed further evidence of clonality within each ST group: all ST101 were group B1, ST405 and ST648 were group D and ST410 were group A. Based on these analyses, we focused the remainder of our study on characterisation of the seven *bla*_NDM-1_-positive ST101 strains.

#### The *bla*_CTX-M-15_ gene has been acquired and mobilised by IS*Ecp1* within Clade 1.

MS6207 and MS6201 carry the IS*Ecp1*-*bla*_CTX-M-15_ element in the same chromosomal location within a composite transposon (Tn*6601*) (Supplementary Table S8) that is inserted in the equivalent location as RD12 in MS6192 and MS6193. In MS6203, an IS*Ecp1*-*bla*_CTX-M-15_ element has instead integrated within the *bsc* operon involved in cellulose biosynthesis, splitting *bscC*. MS6192, MS6193 and MS6194 also carry a chromosomally integrated IS*Ecp1*-*bla*_CTX-M-15_ element, however, in these cases IS*Ecp1*-*bla*_CTX-M-15_ is encoded within a larger transposon that includes a partial Tn*3*, IS*26* and a partial IS from the IS*3* family. This *bla*_CTX-M-15_ transposon is located upstream of the *glg* operon involved in glycogen metabolism. MS6204 encodes two copies of *bla*_CTX-M-15_ on two different plasmids (IncI1 and F-type). The first CTX-M-15 element is comprised of a 99 bp fragment of IS*Ecp1* upstream of *bla*_CTX-M-15_ and is located within a Tn*2* on an Incl1 plasmid backbone, homologous to the CTX-M-15 element of the *E. coli* pJIEi13 plasmid (Genbank accession: EU418923). The second CTX-M-15 element is comprised of a 674bp fragment of IS*Ecp1* that has been truncated at the 5’ end by the insertion of an IS*26*, immediately upstream of *bla*_CTX-M-15_, with the surrounding region punctuated by IS insertions and resistance gene modules within the F-type plasmid pMS6204B, which has the pIP1206-like plasmid backbone.

#### The seven PacBio sequenced NDM-positive ST101 strains are multidrug resistant.

We tested the AMR phenotype of the seven PacBio sequenced ST101 strains using broth microdilution for a total of 35 drugs in more than nine different antimicrobial classes. Based upon the European Committee on Antimicrobial Susceptibility Testing (EUCAST) breakpoint tables v9.0 for the MIC presented, only tigecycline and colistin retained activity against all strains tested (Supplementary Dataset, Table S9). Overall, these isolates are extensively multidrug resistant, particularly to penicillins, cephalosporins, fluoroquinolones and aminoglycosides, confirming our genotyping. Interestingly, all seven strains were resistant to the carbapenems: meropenem, doripenem, ertapenem and imipenem except MS6207, which was resistant to doripenem and ertapenem, but was categorised with intermediate susceptibility to meropenem and imipenem. Carbapenem MIC differences have previously been observed in ST101 strains harbouring similar AMR gene repertoires (17).

#### Deletions in the IncC plasmid backbone alter conjugation efficiency in two *E. coli* ST101 strains.

Although pMS6201A-NDM, pMS6203A-NDM and pMS6207A-NDM share a large IncC-plasmid backbone, in pMS6201A-NDM this region is punctuated with numerous large insertions and deletions. The most notable of these insertions is the previously described Tn*3000*-*bla_NDM-1_* composite transposon, absent in both pMS6203A-NDM and pMS6207A-NDM. Deletions in pMS6201A (as compared to pMS6207A and pMS6203A) include a 12,178 bp region downstream of the *parAB* locus containing hypothetical proteins, a putative ^5m^C MTase, DNA topoisomerase III (*topB*) and a 1,406 bp fragment of *traI*. Downstream of the *bla*_CMY_ gene, pMS6207A-NDM contains a large 22,585 bp deletion in comparison to pMS6203A-NDM, which contains several conjugative transfer genes including, *traC*, *trhF*, *traW*, *traU* and *traN*. A similar, but smaller, deletion of 11,711 bp was observed at the same locus in pMS6201A-NDM containing *traW*, *traU*, *traN* and a 70 bp fragment of *trhF*. Plasmid pMS6203A-NDM is the only IncC plasmid to retain the full composition of conjugation genes, suggesting that the plasmids pMS6201A-NDM and pMS6207A-NDM may be non-conjugative and thus only capable of vertical transmission. To confirm this, we attempted to transfer plasmids by conjugation from MS6201, MS6203 and MS6207 to the *E. coli* J53 strain (8, 9), with only MS6203 generating any transconjugants (6.39 (± 4.28) x 10^-2^).

#### IncC ARI-A regions are highly variable within IncC plasmids pMS6201A-NDM, pMS6203-NDM and pMS6207-NDM.

There are large structural differences within the ARI-A of IncC plasmids pMS6201A-NDM, pMS6203-NDM and pMS6207-NDM. For example, in pMS6207A-NDM, ARI-A encompasses 2 class 1 integrons (In177 and In27) encoding *bla*_OXA-2_ (beta lactam resistance) or *dfrA12* (trimethoprim resistance) and *aadA2* (aminoglycoside resistance) respectively, as well as the ∆IS*Aba125*-*bla*_NDM-1_ module. In pMS6203A-NDM, In27 is missing, with this carrying the *aac(3)-lld* and *tmrB* genes encoding aminoglycoside and tunicamycin resistance respectively, flanked by an IS*Aba14* and an IS*Cfr1*. In pMS6201A-NDM, only In27 is present, however it has integrated into a similar plasmid location as In177 in pMS6207A-NDM, immediately downstream of Tn*6196*, with the ∆IS*Aba125*-*bla*_NDM-1_ module encoded within the IncC plasmid backbone on Tn*3000*.

### Supplementary Figures.

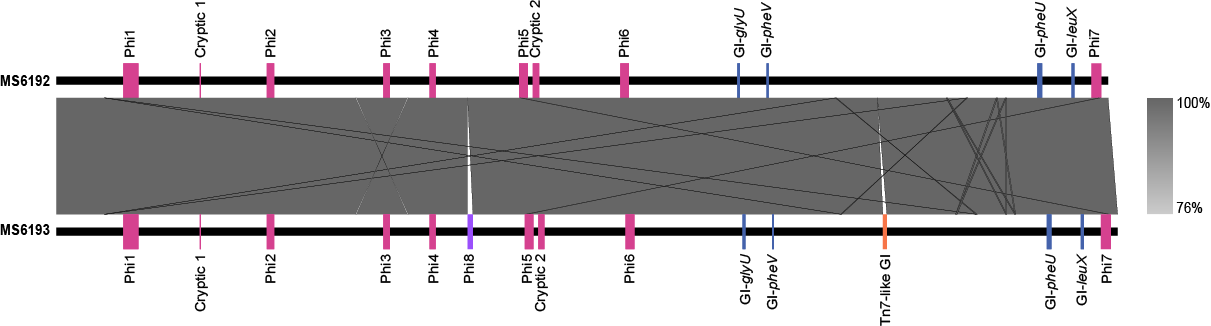

#### Fig S1. Whole genome nucleotide pairwise comparisons between *E. coli* ST101 strains MS6192 and MS6193.

Linear nucleotide alignment of the whole genome, highlighting the 2 extra MGEs in the MS6193 chromosome. Top, *E. coli* MS6192; Bottom, *E. coli* MS6193. Grey shading indicates nucleotide identity between sequences according to BLASTn (76-100%). Shared Prophages and Genomic Islands are shown in pink and blue, respectively. Prophages and genomic islands different to those in *E. coli* MS6192 are shown in purple and orange, respectively. Image prepared using EasyFig.

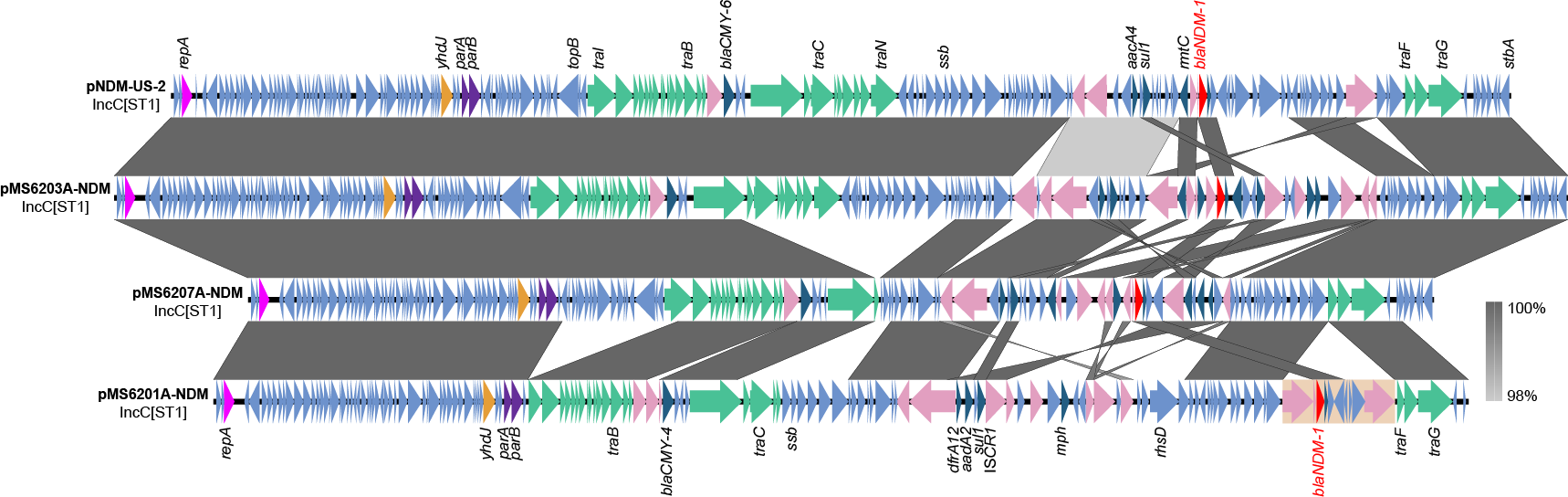

#### Fig S2. Comparisons of *E. coli* ST101 Clade 1 IncC plasmids with pNDM-US-2.

Schematic diagram illustrating the genetic organisation and conserved backbone of NDM+ IncC plasmids. The pNDM-US-2 plasmid (Genbank accession: KJ588779) was included for comparison. Grey shading indicates nucleotide identity between sequences according to BLASTn (98-100%). Key genomic features are indicated, including replication genes: dark pink, maintenance genes: purple, conjugation genes: light green, MTases: orange, IS/Tns: light pink, AMR genes: teal, other CDSs: light blue. The Tn*3000* composite transposon is indicated by an orange rectangle. The *bla*_NDM-1_ gene is labelled in red. Image created using EasyFig.

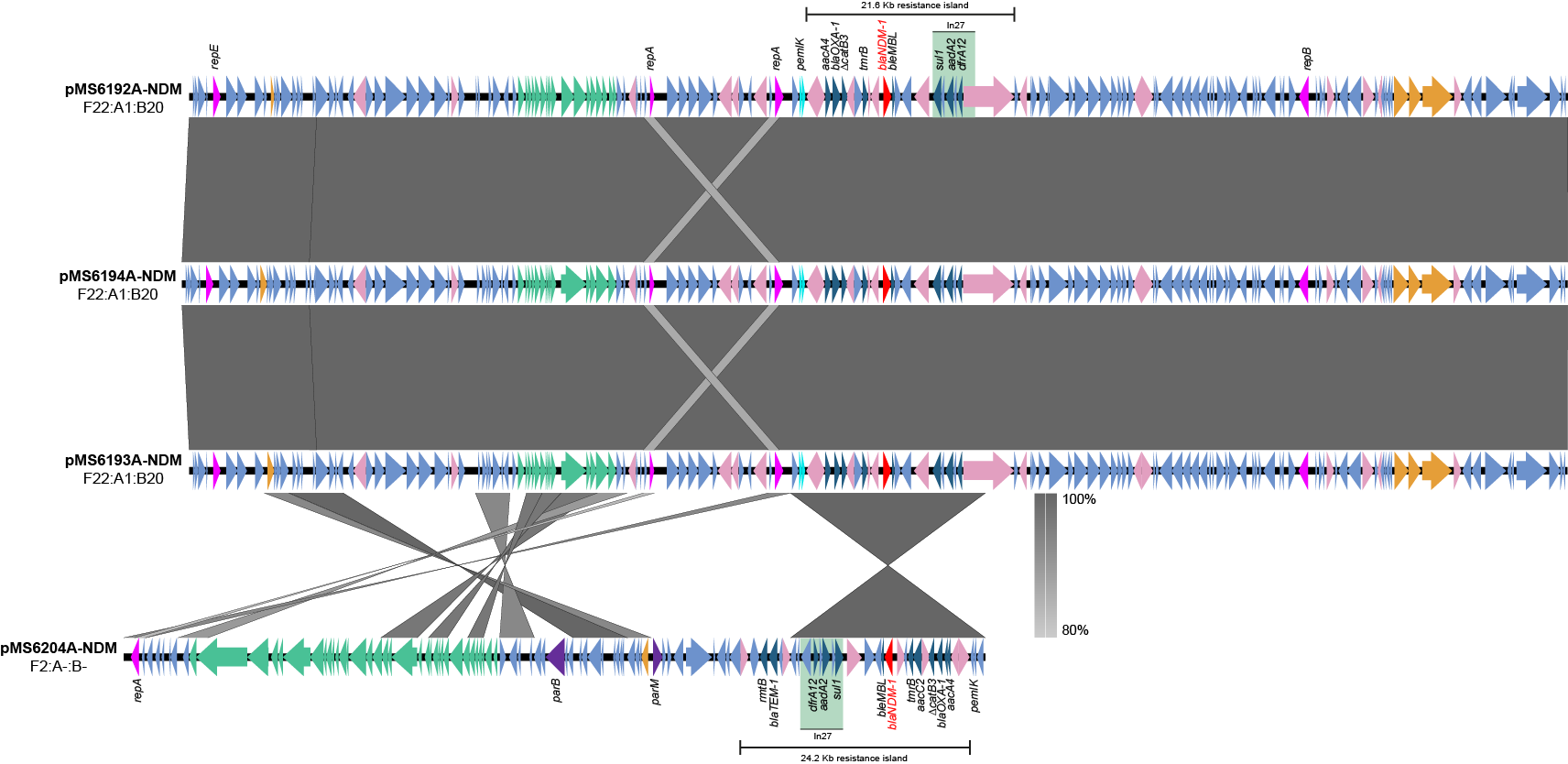

#### Fig S3. Comparisons of NDM-1-positive F-type plasmids from *E. coli* ST101 Clade 1 strains.

Schematic diagram illustrating the genetic organisation of multidrug resistance regions and conserved plasmid backbones. These plasmids feature a number of integrons, IS and transposons which mediate the differing multidrug resistance genotypes. The resistance islands contain the class 1 integron (In27) as well as numerous antimicrobial resistance genes as indicated. Grey shading indicates nucleotide identity between sequences according to BLASTn (80-100%). Key genomic features are indicated, including replication genes: dark pink, maintenance genes: purple, conjugation genes: light green, restriction modification systems and MTases: orange, IS/Tns: light pink, AMR genes: teal, *pemIK* operon: aqua, other CDSs: light blue. Integrons: light green rectangles. The *bla*_NDM-1_ gene is labelled in red. Image created using EasyFig.

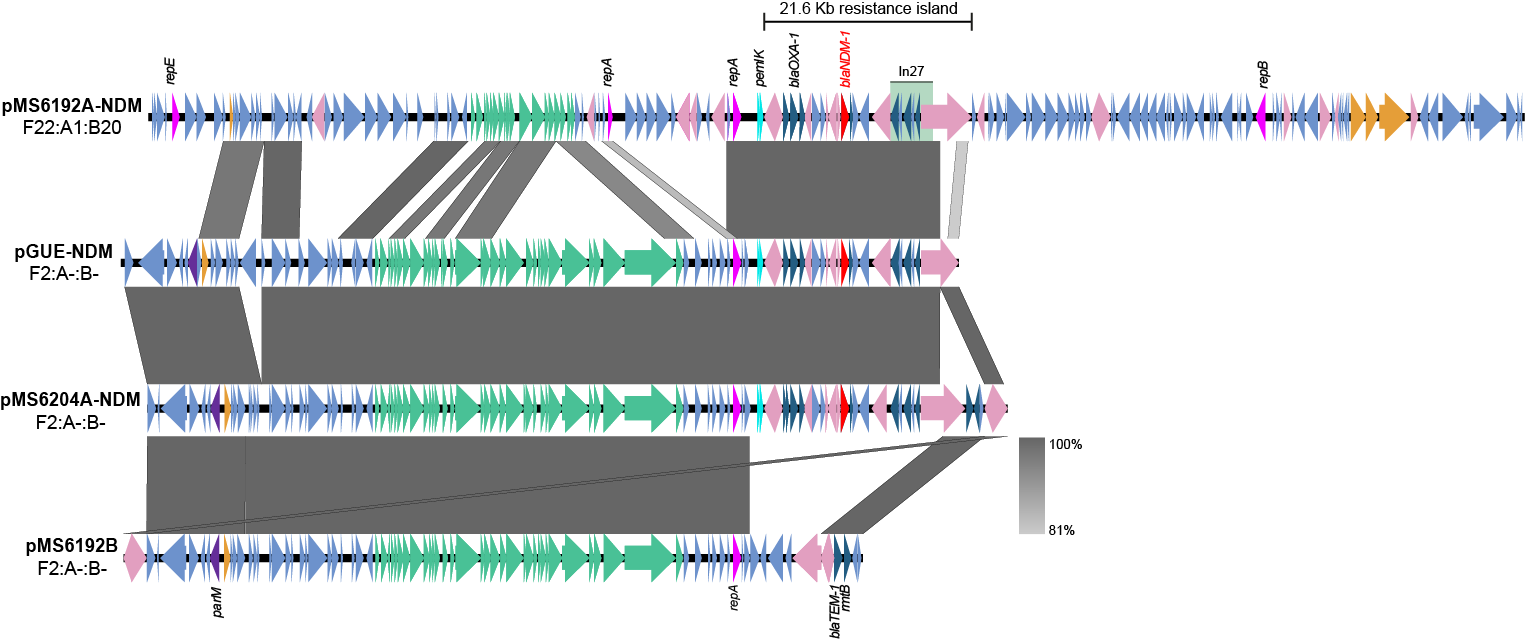

#### Fig S4. Comparisons of plasmid types with a focus on the F2:A-:B- pGUE-NDM backbone and the *bla*_NDM-1_ resistance island.

Nucleotide comparisons between the F-type pIP1206-like plasmid pMS6192A-NDM, the reference plasmid pGUE-NDM (Genbank accession: JQ364967), pMS6204A-NDM and pMS6192B highlighting the shared resistance island and plasmid backbones. pMS6204A and pGUE-NDM have been reverse complemented for easier visualisation. Grey shading indicates nucleotide identity between sequences according to BLASTn (81-100%). Key genomic regions are indicated: IS/Tns: pink, replication genes: dark pink, maintenance genes: purple, conjugation genes: light green, restriction modification systems and MTases: orange, *pemIK* operon: aqua, AMR genes: teal, other CDSs: light blue. Class 1 integrons are shown with a light green rectangle. The *bla*_NDM-1_ gene is labelled in red. Image created using EasyFig.

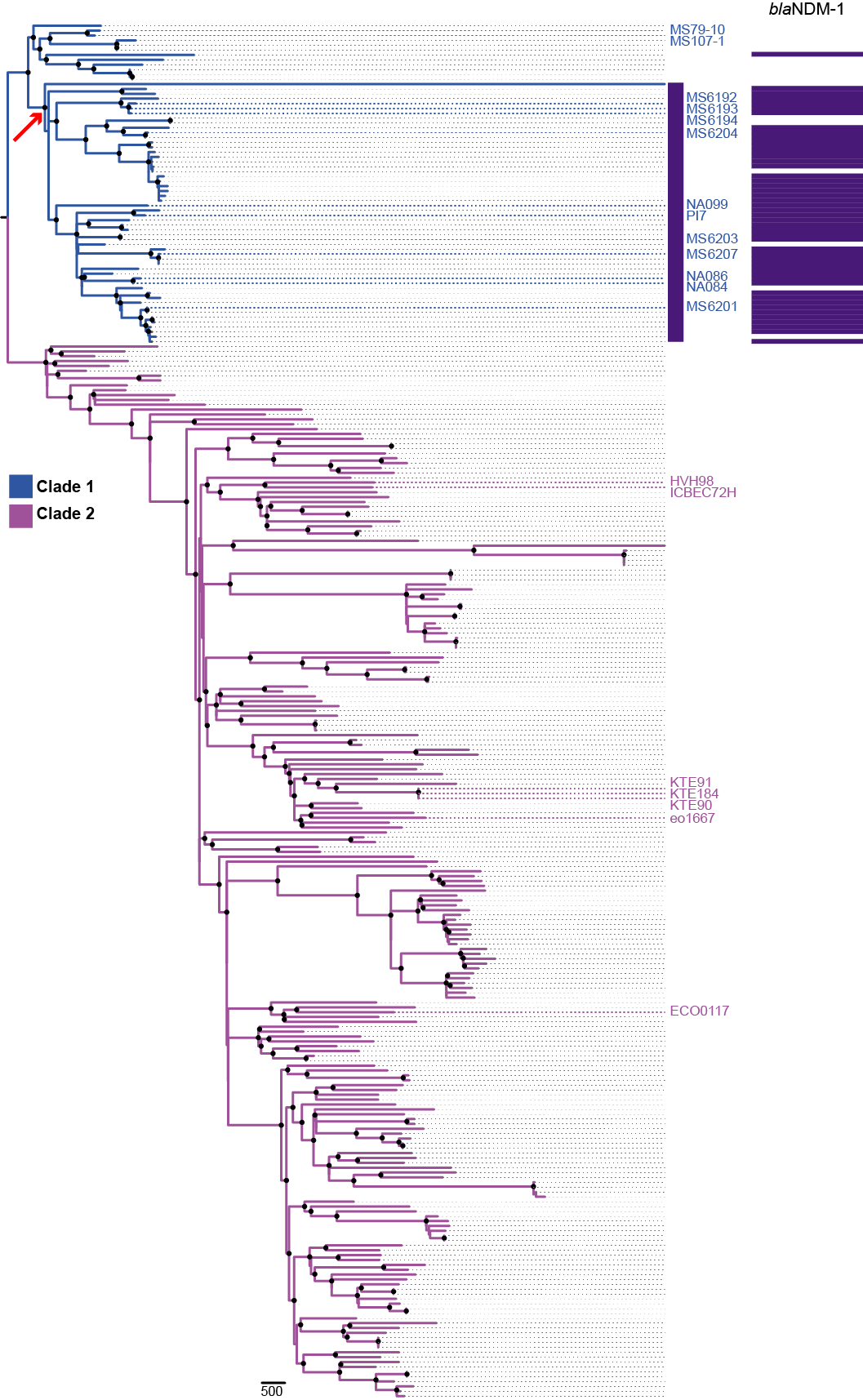

#### Fig S5. Maximum Likelihood phylogenetic relationship of 283 *E. coli* ST101 genomes.

A mid-point rooted, recombination-filtered phylogram was built from 14,185 core-genome SNPs. Taxa labels are coloured by Clade. Clade 1: blue, Clade 2: pink. The 20 *E. coli* ST101 genomes included in this study are highlighted and labelled. Labels of adjacent taxa (e.g. MS6192, MS6193 and MS6194) are spaced out for clarity. 90-100% bootstrap support (1000 replicates) indicated by black nodes. Presence (purple) of *bla*_NDM-1_ (or its variants) was determined by BLASTn (>=95% ID, >60% query coverage) and converted to a presence/absence matrix for input into Phandango (18). Red arrow indicates the branch at which the acquisition of *bla*_NDM-1_ into Clade 1 occurred. Strain names and order are available in Supplementary Dataset 1, Table S13.­

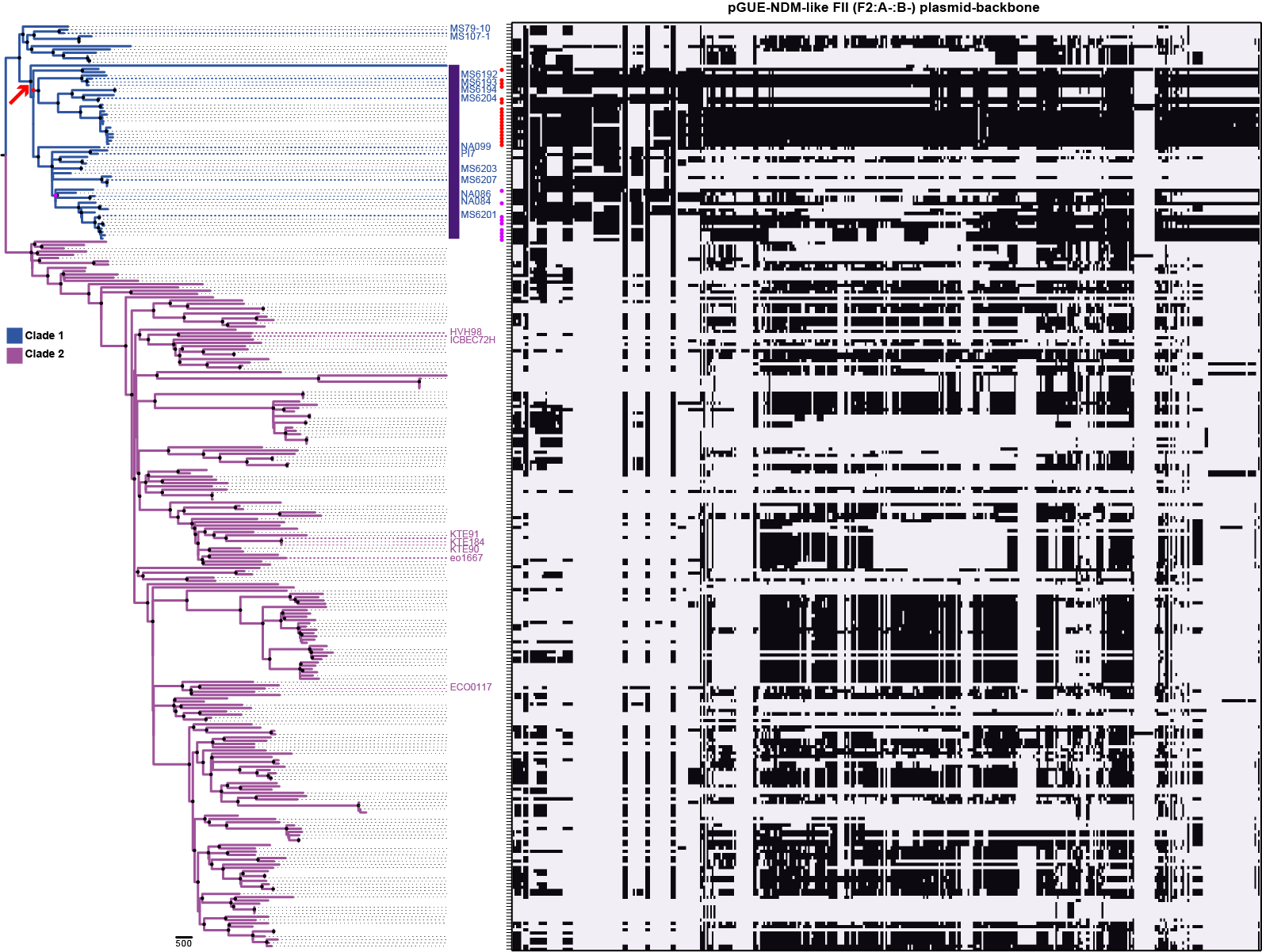

#### Fig S6. Conservation of the pGUE-NDM-like FII (F2:A-:B-) plasmid backbone in 283 *E. coli* ST101 genomes.

Black shading indicates a match of ≥95% nucleotide identity in minimum windows of 200 bp segments, calculated by comparing the query sequence to the assembled contigs for each strain, as implemented in SeqFindr (<https://github.com/BeatsonLab-MicrobialGenomics/SeqFindR>). ST101 strains are ordered according to the phylogenetic relationship defined in Fig S5. Red arrow indicates the branch at which the acquisition of *bla*_NDM-1_ into Clade 1 occurred. Red dots indicate strains that contain a mostly full-length version of the plasmid backbone. Pink dots indicate strains that contain plasmids with homology to several pGUE-NDM plasmid modules.

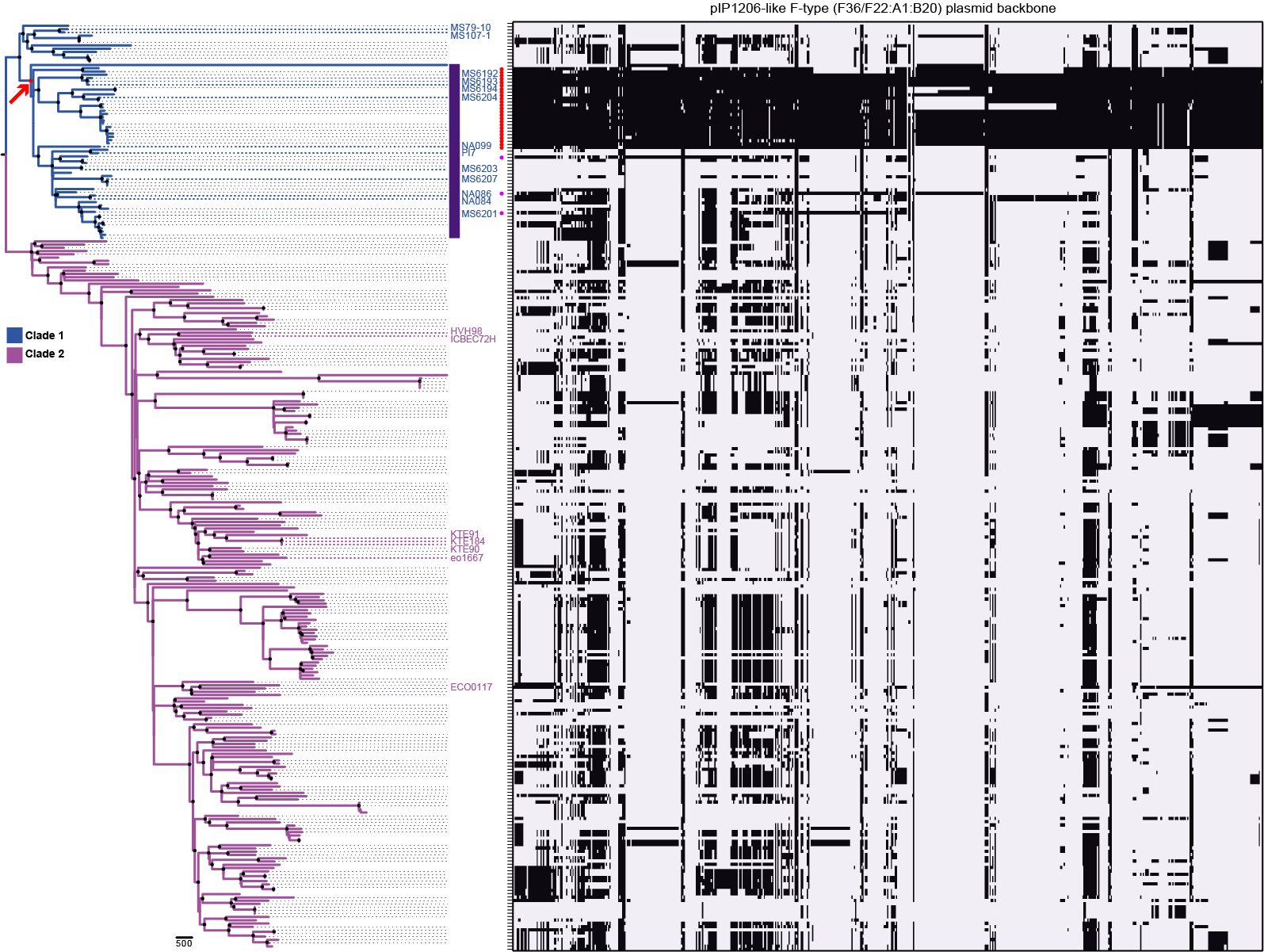

#### Fig S7. Conservation of the pIP1206-like F-type (F36/F22:A1:B20) plasmid backbone in 283 *E. coli* ST101 genomes.

Black shading indicates a match of ≥95% nucleotide identity in minimum windows of 200 bp segments, calculated by comparing the query sequence to the assembled contigs for each strain, as implemented in SeqFindr. ST101 strains are ordered according to the phylogenetic relationship defined in Fig S5. Red arrow indicates the branch at which the acquisition of *bla*_NDM-1_ into Clade 1 occurred. Red dots indicate strains that contain a mostly full-length version of the plasmid backbone. Pink dots indicate strains that contain plasmids with homology to several pIP1206 plasmid modules.

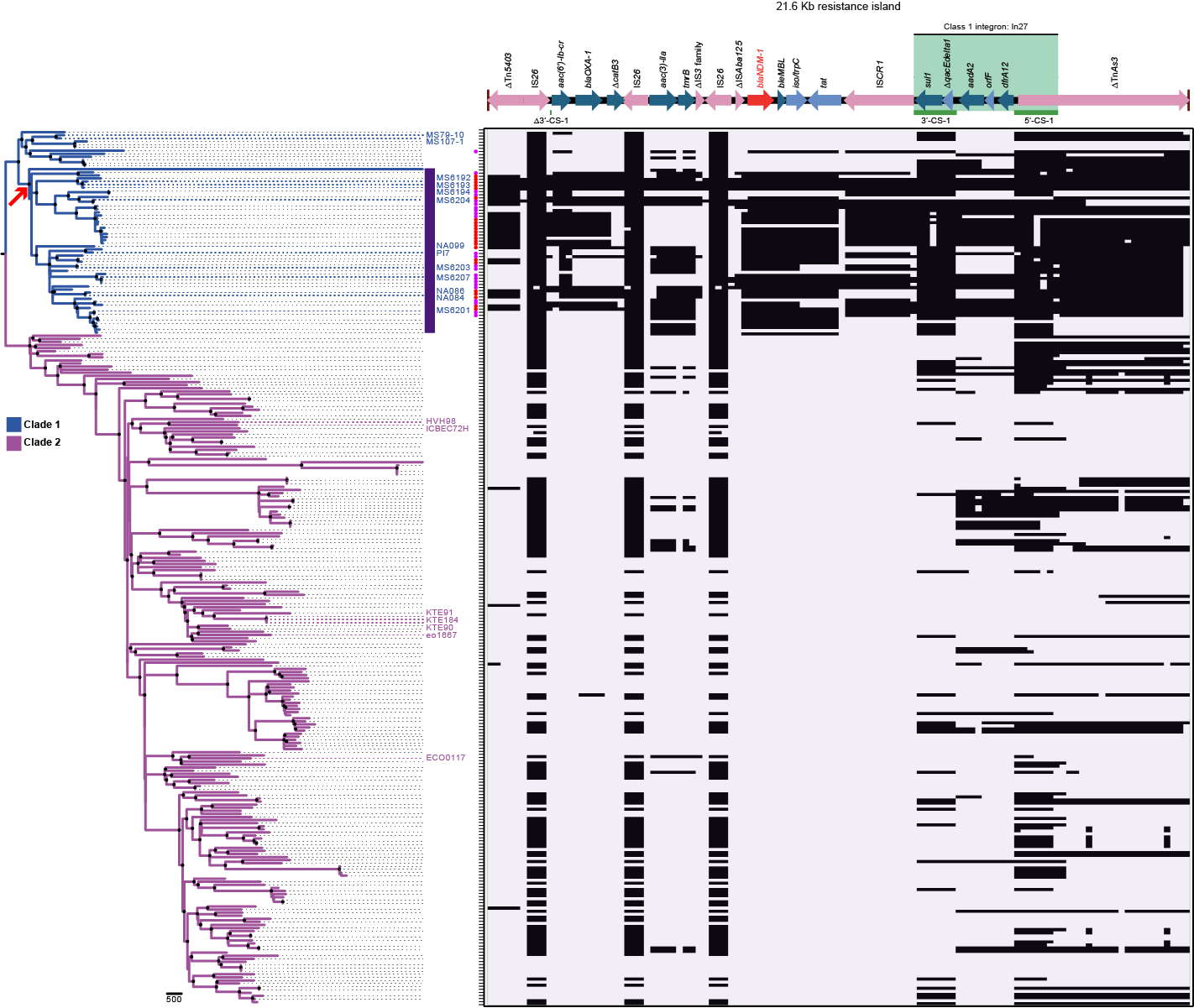

#### Fig S8. Conservation of the *bla*_NDM-1_ resistance island in 283 *E. coli* ST101 genomes.

Black shading indicates a match of ≥95% nucleotide identity in minimum windows of 200 bp segments, calculated by comparing the query sequence to the assembled contigs for each strain, as implemented in SeqFindr. ST101 strains are ordered according to the phylogenetic relationship defined in Fig S5. Red arrow indicates the branch at which the acquisition of *bla*_NDM-1_ into Clade 1 occurred. Red dots indicate strains that contain a mostly full-length version of the resistance island with both flanking transposons. Pink dots indicate strains that contain homology to several resistance island modules including at least one flanking transposon.

### References

1.	Djoko KY, Achard MES, Phan MD, Lo AW, Miraula M, Prombhul S, Hancock SJ, Peters KM, Sidjabat H, Harris PN, Mitic N, Walsh TR, Anderson GJ, Shafer WM, Paterson DL, Schenk G, McEwan AG, Schembri MA. 2017. Copper ions and coordination complexes as novel carbapenem adjuvants. Antimicrob Agents Chemother doi:10.1128/aac.02280-17.

2.	Kumarasamy KK, Toleman MA, Walsh TR, Bagaria J, Butt F, Balakrishnan R, Chaudhary U, Doumith M, Giske CG, Irfan S, Krishnan P, Kumar AV, Maharjan S, Mushtaq S, Noorie T, Paterson DL, Pearson A, Perry C, Pike R, Rao B, Ray U, Sarma JB, Sharma M, Sheridan E, Thirunarayan MA, Turton J, Upadhyay S, Warner M, Welfare W, Livermore DM, Woodford N. 2010. Emergence of a new antibiotic resistance mechanism in India, Pakistan, and the UK: a molecular, biological, and epidemiological study. Lancet Infect Dis 10:597-602.

3.	Clermont O, Bonacorsi S, Bingen E. 2000. Rapid and Simple Determination of the Escherichia coli Phylogenetic Group. Applied and Environmental Microbiology 66:4555-4558.

4.	Wirth T, Falush D, Lan R, Colles F, Mensa P, Wieler LH, Karch H, Reeves PR, Maiden MC, Ochman H, Achtman M. 2006. Sex and virulence in Escherichia coli: an evolutionary perspective. Mol Microbiol 60:1136-51.

5.	Johnson JR, Stell AL. 2000. Extended virulence genotypes of Escherichia coli strains from patients with urosepsis in relation to phylogeny and host compromise. J Infect Dis 181:261-72.

6.	Poirel L, Dortet L, Bernabeu S, Nordmann P. 2011. Genetic features of blaNDM-1-positive Enterobacteriaceae. Antimicrob Agents Chemother 55:5403-7.

7.	Hancock SJ, Phan MD, Peters KM, Forde BM, Chong TM, Yin WF, Chan KG, Paterson DL, Walsh TR, Beatson SA, Schembri MA. 2017. Identification of IncA/C Plasmid Replication and Maintenance Genes and Development of a Plasmid Multilocus Sequence Typing Scheme. Antimicrob Agents Chemother 61.

8.	Yi H, Cho Y-J, Yong D, Chun J. 2012. Genome Sequence of Escherichia coli J53, a Reference Strain for Genetic Studies. Journal of Bacteriology 194:3742-3743.

9.	Jacoby GA, Han P. 1996. Detection of extended-spectrum beta-lactamases in clinical isolates of Klebsiella pneumoniae and Escherichia coli. Journal of Clinical Microbiology 34:908-911.

10.	Li H. 2018. Minimap2: pairwise alignment for nucleotide sequences. Bioinformatics 34:3094-3100.

11.	Li H, Durbin R. 2009. Fast and accurate short read alignment with Burrows-Wheeler transform. Bioinformatics 25:1754-60.

12.	Li H, Handsaker B, Wysoker A, Fennell T, Ruan J, Homer N, Marth G, Abecasis G, Durbin R. 2009. The Sequence Alignment/Map format and SAMtools. Bioinformatics 25:2078-9.

13.	Quinlan AR, Hall IM. 2010. BEDTools: a flexible suite of utilities for comparing genomic features. Bioinformatics 26:841-2.

14.	Kolmogorov M, Yuan J, Lin Y, Pevzner PA. 2019. Assembly of long, error-prone reads using repeat graphs. Nat Biotechnol 37:540-546.

15.	Carver T, Harris SR, Berriman M, Parkhill J, McQuillan JA. 2012. Artemis: an integrated platform for visualization and analysis of high-throughput sequence-based experimental data. Bioinformatics 28:464-9.

16.	Carver TJ, Rutherford KM, Berriman M, Rajandream MA, Barrell BG, Parkhill J. 2005. ACT: the Artemis Comparison Tool. Bioinformatics 21:3422-3.

17.	Yoo JS, Kim HM, Koo HS, Yang JW, Yoo JI, Kim HS, Park HK, Lee YS. 2013. Nosocomial transmission of NDM-1-producing Escherichia coli ST101 in a Korean hospital. J Antimicrob Chemother 68:2170-2.

18.	Hadfield J, Croucher NJ, Goater RJ, Abudahab K, Aanensen DM, Harris SR. 2018. Phandango: an interactive viewer for bacterial population genomics. Bioinformatics 34:292-293.
